# Pathologies of Between-Groups Principal Components Analysis in Geometric Morphometrics

**DOI:** 10.1101/627448

**Authors:** Fred L. Bookstein

## Abstract

Good empirical applications of geometric morphometrics (GMM) typically involve several times more variables than specimens, a situation the statistician refers to as “high *p/n*,” where *p* is the count of variables and *n* the count of specimens. This note calls your attention to two predictable catastrophic failures of one particular multivariate statistical technique, between-groups principal components analysis (bgPCA), in this high-*p/n* setting. The more obvious pathology is this: when applied to the patternless (null) model of *p* identically distributed Gaussians over groups of the same size, both bgPCA and its algebraic equivalent, partial least squares (PLS) analysis against group, necessarily generate the appearance of huge equilateral group separations that are actually fictitious (absent from the statistical model). When specimen counts by group vary greatly or when any group includes fewer than about ten specimens, an even worse failure of the technique obtains: the smaller the group, the more likely a bgPCA is to fictitiously identify that group as the end-member of one of its derived axes. For these two reasons, when used in GMM and other high-*p/n* settings the bgPCA method very often leads to invalid or insecure bioscientific inferences. This paper demonstrates and quantifies these and other pathological outcomes both for patternless models and for models with one or two valid factors, then offers suggestions for how GMM practitioners should protect themselves against the consequences for inference of these lamentably predictable misrepresentations. The bgPCA method should never be used unskeptically — it is never authoritative — and whenever it appears in partial support of any biological inference it must be accompanied by a wide range of diagnostic plots and other challenges, many of which are presented here for the first time.

## 1. Introduction

Figure 1 shows two scatterplots based on analyses of the same simulated data set of 30 specimens divided into three groups of ten each. The specimens are modeled as having been “measured” on a total of 300 variables, for instance the Procrustes shape space for 152 landmarks and semilandmarks in two dimensions, but the distribution I am simulating for these 300 dimensions is the most uninformative possible: 300 independent Gaussian (normal) variables of the same mean (here, zero) and the same variance (here, 1.0) for each of the thirty simulated cases, which have been grouped entirely arbitrarily (the first ten specimens, the second ten, the third ten). The scatterplot on the right is the usual display from a Partial Least Squares (PLS) analysis of group against the 300-variable observation vector; that on the left is the similarly conventional display from a “between-group principal components analysis” (bgPCA) of the derived sample of three group means in the same 300-dimensional space. In either panel, the printed symbol corresponds to the imputed group index, which is 1, 2, or 3. Keep in mind that these groups are arbitrary subsamples of simulated specimens that were in fact identically distributed independent of group.

**Figure 1.**
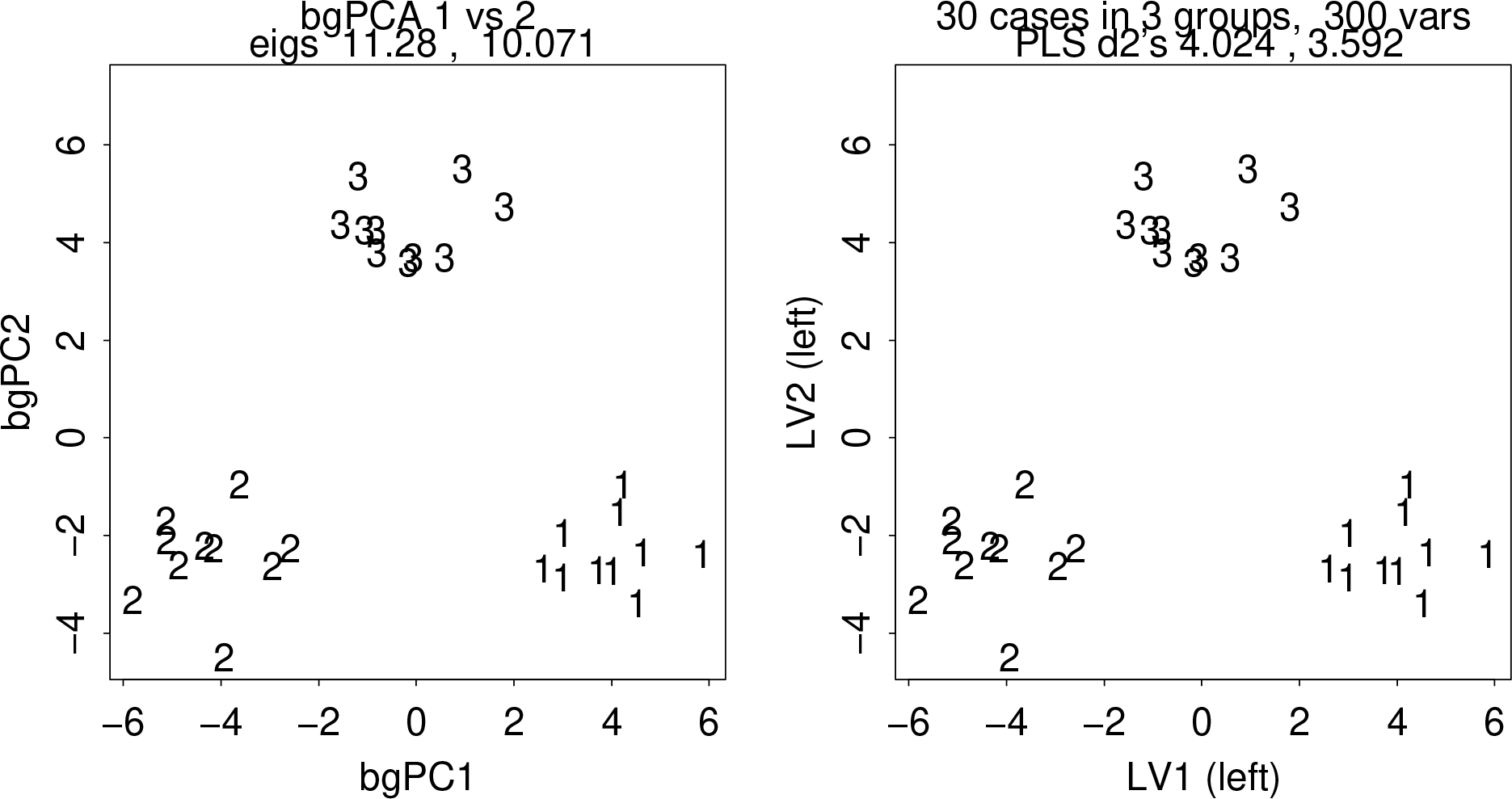
A typical manifestation of the pathology that is the topic of this note: the completely fictitious production of group separations when either of two equivalent techniques — bgPCA or PLS against a group dummy array — is applied to the same completely uninformative data set of 300 identically distributed Gaussian variables on 30 simulated “specimens.” Plotting characters 1, 2, 3 correspond to the (entirely arbitrary) group index assigned to the thirty “specimens” in three sets of ten. The group separation is indeed startling and would be assessed as hugely significant by any statistical maneuver that was unaware of the ordination’s actual origin in a data set of 300 variables identically distributed over 30 specimens. Thus to infer a group separation from this shared scatterplot is clearly a mistake.

These are evidently the same scatterplot. Furthermore, they are described by equivalent figures of merit — the quantities listed in their subtitles. For bgPCA, on the left, these are the eigenvalues (net variance explained) of the only two principal components of those three group means; each is a component of the total sum of squares over all the variables, which is about 300. For PLS, on the right, these are the total “squared covariances explained” between the same vector of 300 simulated measurements and a 3-by-30 matrix of group dummy variables each of mean 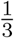 and variance 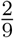. The two pairs of entries have the same ratio, about 1.12 to 1, and their ratio of 2.803 from right to left is precisely 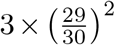, the count of groups times the undoing of the factor that my statistical software package applies when it computes variances. That bgPCA and PLS against a group dummy are algebraically the same has been known for a while (see, e.g., Boulesteix, 2005). The figures to follow will invoke whichever one offers the simpler explanation of the pathology that is my main topic in this section.

That pathology, already evident in Figure 1, can be quantified by an application to this small-group setting of the celebrated *Marchenko-Pastur theorem* (MPT) that I reviewed in a previous publication in this Journal (Bookstein, 2017). The MPT sets out a formula for the limiting distribution of *all* the eigenvalues of a data set of *p* identically distributed standard Gaussian variables (mean 0, variance 1) on *n* specimens as both numbers tend to infinity in a fixed ratio *y* = *p/n*. The theorem states a limit for the maximum of these eigenvalues, 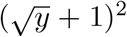, and a similar-looking limit 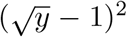 for their minimum. These formulas apply “asymptotically,” as the statisticians say. The analysis in Section 2 will show that to apply the formula to the scenario in Figure 1 we substitute the value 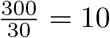 for *y* and multiply by 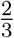 (for the case of just three groups), resulting in a value of 11.55, quite close to the observed eigenvalue of 11.28 printed in the left panel of the figure. The sum of the two numbers there is 21.35, whereas the total variance of the three group mean 300-vectors is expected to be 20 (see, again, Section 2), likewise a good match.

Then the essence of this first pathology leaps to the eye here. While the variances of the group means themselves total about 20, the variance of any direction *within* a group remains near the value of 1.0 that characterizes almost every projection of this 300-dimensional scatter onto a line. So the group means *must* be separated by about four times their within-group standard deviation; in other words, the scatters must be perfectly separated by group. But there is no such separation in the model we are simulating — the groups were drawn from *identical* high-dimensional distributions. The separations were produced *solely* by the bgPCA machinery itself. This seems like a serious, indeed fatal, mathematical error for any technique that is ever employed in a context of ordination.

Some hints in this figure will concern us in the sequel. For instance, the triangle of the centroids of the three groups in the figure appears to be oddly close to equilateral. We can check this hunch by repeating the simulation a dozen times. As Figure 2 shows, we always get *precisely* the same “answer”: three groups cleanly separated with centroids forming a nearly equilateral triangle, larger eigenvalue usually 10.0 or a bit above, the pair of them always totalling about 20, and the within-group variances still hovering around 1.0. The equilateral triangle of centroids appears to spin on the page because inasmuch as the eigenvalues of such a shape are always nearly equal, the computed principal axes will be uniformly distributed with respect to the centroids that were their data. And the triangle’s orientation (clockwise or counterclockwise) is unstable because PCA does not constrain the signs of the extracted components.

**Figure 2.**
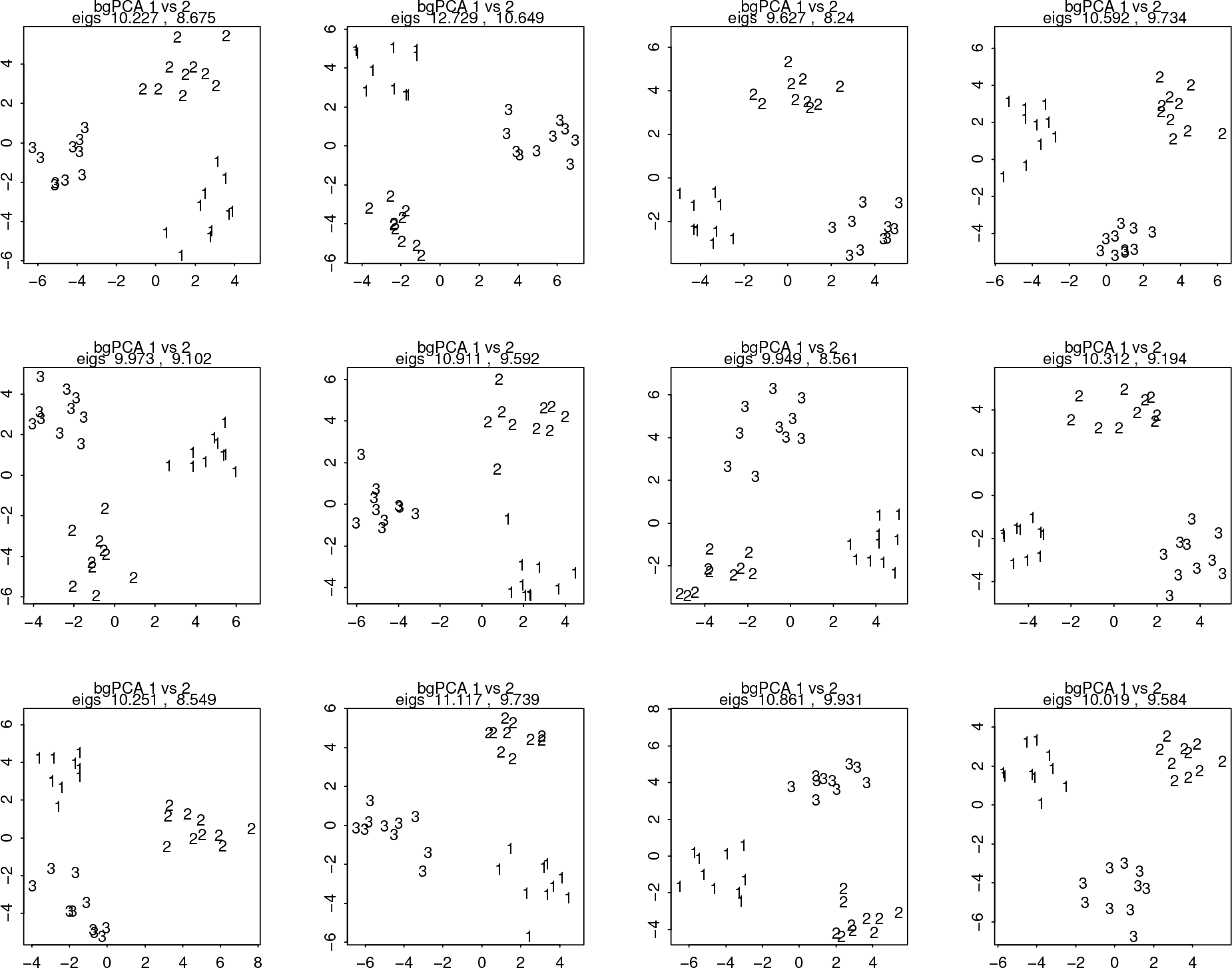
Twelve replications of the simulation in Figure 1. The result is always a separation of the “groups” with eigenvalues of the bgPCA analysis nearly equal, their total always about 20, and within-group variances always about 1.0; hence, always, the fiction of perfect separation.

Let us explore a little further. Figure 3 presents the same analysis as in Figure 1 but for four groups of ten specimens each, instead of three, and a total of count of variables likewise multiplied by 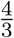, to 400, so that the value *y* = *p/n* remains at 10. Now we see a large-scale quadrilateral of group-specific scatters, still with within-group variances around 1 and still perfectly separated but now no longer quite so symmetrically placed on the plot. But that is because we have projected down onto only two dimensions. When there are four groups the space of the bgPC’s is actually three-dimensional, as in Figure 4, and the third eigenvalue is not negligible with respect to the first two. When we plot all three of these dimensions, we see that indeed the group centroids fall at the vertices of an equilateral *tetrahedron*, and the eigenvalues remain very nearly equal and total approximately 30, which is the trace of the expected variance-covariance matrix of the corresponding quartet of group means.

**Figure 3.**
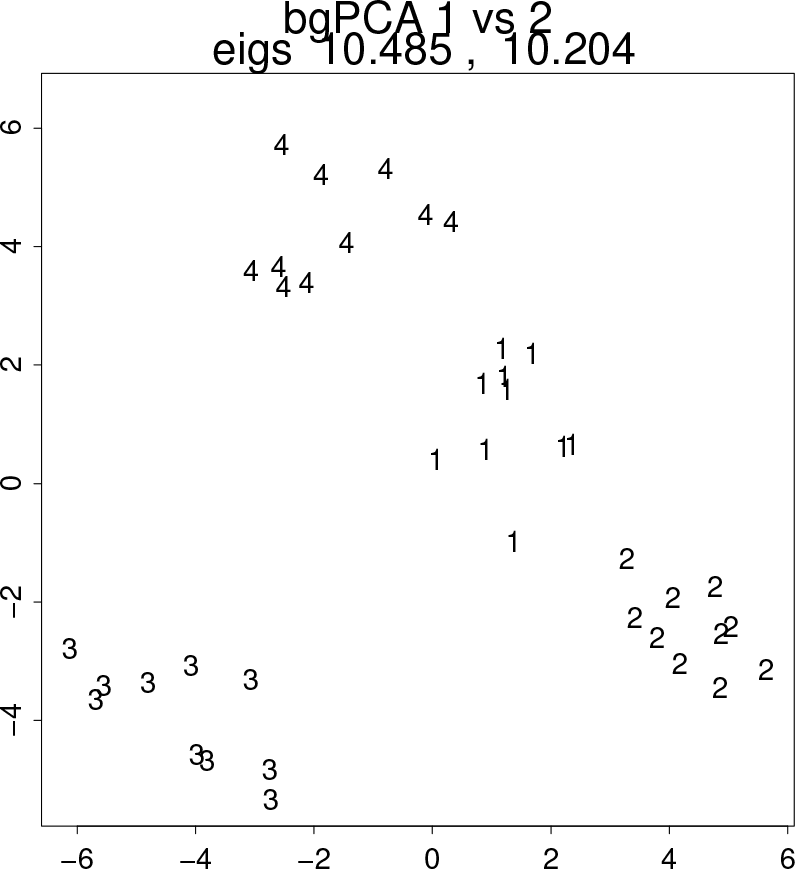
Analogous simulation for four groups of ten specimens each and likewise ten times as many variables as specimens, now a total of 400. The techniques of bgPCA and PLS are still identical, and likewise the fictitious “finding” of perfect separation, but the symmetry of the plots in Figures 1 and 2 appears to be broken.

**Figure 4.**
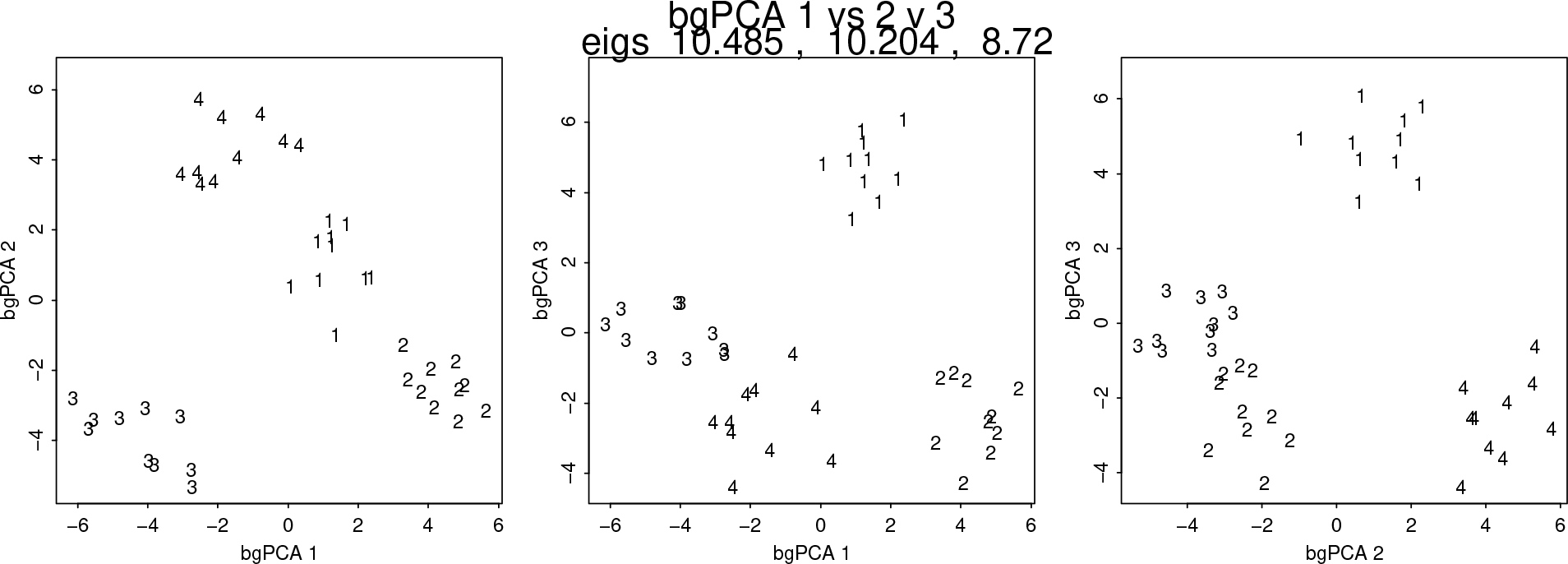
Completion of Figure 3 by including bgPC3, the final between-group principal component. The symmetry of the configuration of four group centroids is restored by a display in three dimensions instead of two — we are looking at three orthogonal but otherwise arbitrary projections of an almost-equilateral tetrahedron.

Similarly we can explore the dependence of this pathology on the ratio *y* = *p/n* of variables to cases. Raising this ratio to 40, for instance, while remaining with four groups of ten specimens each, results in the three-dimensional scatter of Figure 5, which continues to be the display of an equilateral tetrahedron. The three eigenvalues remain nearly equal but now their total is very nearly 120, four times what it was for Figure 4, as there are four times as many simulated variables of unit variance. Back in the setting of just three groups, Figure 6 repeats the analysis of Figure 2 for twice the count of variables, noticing, again, that the equilateral triangle has twice the eigenvalue total and therefore higher group separations (since the within-group variances remain around unity), while the configuration itself continues to appear just to spin (with reflection).

**Figure 5.**
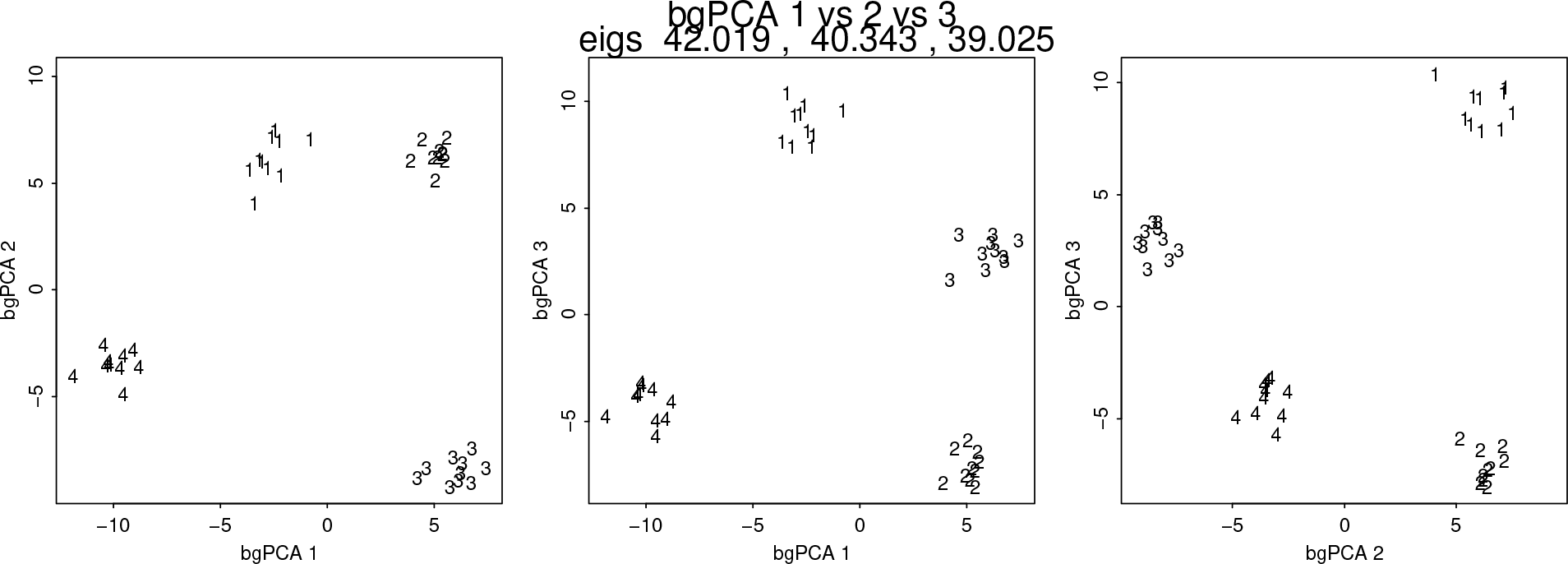
Analogue of Figure 4 with four times as many variables. The group centroids still lie at the vertices of an equilateral tetrahedron, around which variances are still about 1.0 in every direction, while the eigenvalues (always nearly equal) summarizing the separation of the fictitious “group centroids” now total 120 instead of 30.

**Figure 6.**
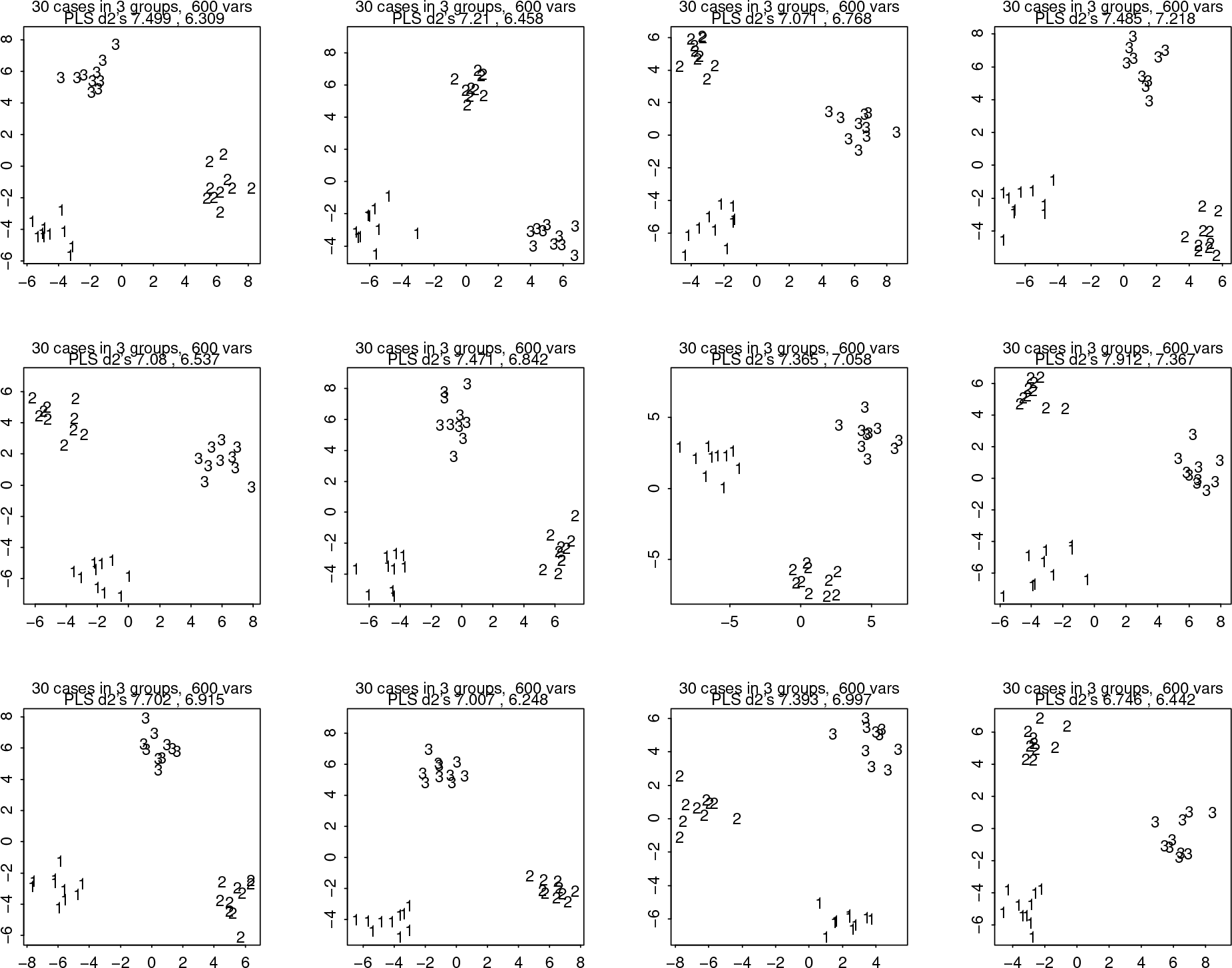
Analogue of Figure 2 for twice as many variables, 600 instead of 300, hence *y* = *p/n* = 20. The configuration of group centroids remains equilateral even as the visual separation of the groups increases in proportion to the square-root of the eigenvalues (here reported as PLS totals of explained squared covariance).

### A brief history

In view of these persistent, potentially devastating problems, I admit with some embarrassment that of the two analytic dialects here, the PLS version seems apparently to be mine, published originally without any citations to precursors in a paper with Randy McIntosh (McIntosh et al., 1996) for which the domain of application was positron emission tomography (PET) imaging. But priority goes to an earlier presentation in the language of bgPCA, Yendle and MacFie (1989),^1^ who named their technique “discriminant principal components analysis” (DPCA) and recommended it when “the number of variables exceeds the number of samples” (i.e., when *p/n* > 1). Section 9.1 of Jolliffe (2002), a standard textbook of PCA, situates the bgPCA approach (via this Yendle–MacFie citation) within a spectrum of variously “prewhitened” versions of canonical variates analysis (CVA). Ironically, even though Yendle and MacFie note that “under certain conditions the discrimination achieved by CVA may be totally spurious,” they do not even hint at the MPT-related problems that are my topic here, which render some discriminations potentially achieved by their DPCA likewise “totally spurious.”

The logic of either presentation, bgPCA or PLS by group, is straightforward: to circumvent the pathologies of a much earlier approach, linear discriminant analysis, in settings where the predictors (for PET, the positron emission densities voxel by voxel; for GMM, the Procrustes coordinates of an arbitrarily long list of landmarks and semilandmarks) are too numerous for their covariance matrix to be invertible, are subject to exact linear constraints that likewise render the covariance matrix noninvertible, or are too intercorrelated for that inverse to be stable over sampling. For a good didactic review of this technique in the context of the difficulties it was intended to circumvent, see Mitteroecker and Bookstein, 2011.

In the days before we biometricians stumbled across the MPT, the error embraced by this biometrical approach was forgivable. It seemed like a good idea to maximize the strength of a grouping signal over linear combinations of measurements, and so we needed a quick, easily programmed, preferably linear fix that permitted interpretation via prior knowledge of within-group factors or selection gradients. None of us carried out simulations extensive enough to recognize either of the fatal flaws examined in this paper. But with the passage of time the conditions conducive to these pathologies have become more common. For instance, the first of these pathologies is the huge instability of PCA and related techniques whenever the ratio *y* of variables to specimens is much larger than 1 and there is substantial commensurate measurement error or other independent variation in most variables separately. But the commensurability of nearly independent measurement errors over variables characterizes the isotropic Mardia–Dryden distribution that lies at the core of many approaches to landmark data analysis in GMM, and with the widespread adoption of the R programming language it is now possible to apply the usual tools of multivariate analysis to hundreds or even thousands of shape coordinates at once. (See Section 4.5 for a workaround.) In many of the fields accustomed to high-*p/n* data sets, wide variation of subgroup sample sizes has become a commonplace, as in recent studies of species of *Homo*, partially bgPCA-driven, by Chen et al. (2019) and Détroit et al. (2019), two papers explicitly critiqued below. Variability of subgroup size when the *p/n* ratio is high is the requisite for this paper’s second main pathology, the automatic alignment of the first one or two bgPCA axes with contrasts of the grand mean against the smallest one or two subgroups only.

For these and other reasons, the frequency with which bgPCA analyses appear in peer-reviewed articles has been increasing even as its underlying flaws remained unknown. A `scholar.google.com` retrieval finds the counts of recent articles that mention bgPCA to be as follows: ^2^ articles published in 2013, 8; in 2014, 13; in 2015, 21; in 2016, 14; in 2017, 21; in 2018, 24; in 2019, through May 10, 14. The present note is a companion piece to Cardini, Rohlf, and O’Higgins (2019), which arrives at the same diagnosis as mine of the first pathology, that of fictitious clustering, somewhat less formally, without invoking the MPT but with several more realistic simulations and simpler algebra. The two approaches, one more biomathematical and one less so, end up offering the same advice: **the technique of bgPCA should never be used with good GMM data**, meaning, data with sufficiently many shape coordinates to represent the form evocatively. That is, bgPCA should never be applied to data sets where the count of shape coordinates is more than a small fraction of the count of specimens. (The recommendation in Mitteroecker and Bookstein 2011 that ignored this restriction of applicability is hereby countermanded by the second of its two authors.) And there is an even deeper flaw, one no longer specific to GMM: the illusion that maximizing variance or covariance over linear combinations of huge numbers of shape coordinates is a reliable source of biological insight.

The purpose of this article is to discuss both of these newly uncovered pathologies of the bgPCA technique, some potential diagnostics, and a range of appropriate responses by any interested disciplinary community. Section 1 has already demonstrated the first of the pathologies, the misrepresentation by unambiguously distinct clusters of high-dimensional Gaussian data that are actually independent of group. Section 2 reviews the underlying pathologies of this paper’s many examples in terms of the governing theorem, the MPT, that accounts for them in quantitative detail. Section 3 shows how these pathologies persist through an enrichment of the null model of totally noninformative Gaussians here to incorporate the factor structures that render the statistical models in organismal applications far more realistic. Section 4 is concerned with tools that permit the evaluation of bgPCA models from the point of view of a principled skeptic or manuscript reviewer. Two standard classes of diagnostic tools, permutation tests and crossvalidation approaches, are rejected; in the course of this second rejection, that second pathology of the bgPCA method is uncovered, the pathology induced by varying subsample counts, which in many paleoanthropological applications is even more catastrophic than the hallucination of clusters. The final Section 5 summarizes the critique by a list of seven pointed recommendations that every user of bgPCA should consider before drawing any scientific inferences from its computations.

## 2. Why are we seeing this?

Where, then, do these formulas come from that the simulations in Section 1 exemplify? The argument in this article and its companion piece is not merely an authoritative collaborative critique of an inadequately tested, insufficiently theorem-driven method prematurely adopted by a few applied communities. Instead it is meant to convey an important new fact about “megavariate data analysis” that is fundamental to the geometry of these data spaces and so might be of interest more generally to evolutionary and developmental biologists who deal with huge data sets like these. The MPT itself is too deep for biologists to apprehend without a struggle, but its application to the bgPCA method is straightforward. Here is how that goes.

Let me standardize notation for this application of the theorem as follows. Set down the bgPCA algorithm in five steps:

1. Collect a data set of *p* variables over *n* specimens divided a priori into *g* subgroups. Assume for simplicity that each of the *g* subgroups has the same count *m* of specimens. (I return to this assumption in Section 4.3.)
2. Compute the mean *p*-vector for each of the subgroups. This gives you *g* new *p*-vectors.
3. Extract the principal components of that set of *g p*-vectors.
4. Impute principal component scores for all *n* of the original specimens using the formulas for the principal components of those *g p*-vectors.
5. Plot the resulting principal component scores, imaginatively interpret the patterns of the first two or three principal component loading vectors or the contrasts implied by the phylogenetics or the experimental design of the groups, and publish or paste onto a poster.

The MPT (Marchenko and Pastur, 1967) is a theorem fundamental to the probability theory of random processes. For an intermittently accessible exposition, see Bookstein, 2017. The theorem states that if *X* is a data matrix of *p* standard Gaussians (mean 0, variance 1) over *n* cases, then in the limit of *p* and *n* both tending to infinity at fixed ratio *y* = *p/n*, the distribution of the nonzero singular values of the *p* × *p* matrix *X*^′^*X/n* — the list of the conventional “explained variances” of the full set of all *p* uncentered principal components of *X* — approaches one specific family of distributions that are a function only of *y*: probability is nonzero only between 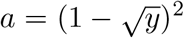 and 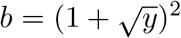 (the notation for *y* < 1), or equivalently between 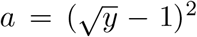 and 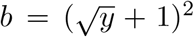 (for other *y*’s), with probability density 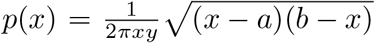 over that interval. The problems I highlighted in Bookstein (2017) arise from the unbounded nature of the ratio of maximum to minimum over this range when *y* is near 1. The present note is concerned only with the higher end of the range, the values 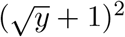 and their transformation into expected bgPCA eigenvalues, for diverse values of *y* mostly much greater than 1.

Even before we examine the way this theorem implied the catastrophic performance of bgPCA as sketched in my first six figures, it is worth sketching its effect on ordinary PCA of null-model simulations similar to those there. As Figure 7 demonstrates, the behavior of PCA under high-*p/n* conditions is not at all what one has been taught to expect. The example is now of 600 cases on 60 independent and identically distributed standard Gaussian variables, hence *y* = *p/n* is still equal to 10. The example no longer concerns any grouping variable; nevertheless the same theorem applies. The upper left panel shows a random sample of 25 pairwise scatters. If the data set were usefully described by explanatory factors, at least some of these distributions would appear to be noncircular ellipses, but here their visual diameter is quite homogeneous and their circularity apparent. (The median absolute correlation among all 600 × 599/2 = 179700 possibilities is only 0.0887.)

At upper right is the conventional scree plot of the eigenvalues from a standard principal component analysis — the explained variances of the 59 nonzero centered principal components here. On the conventional null model for infinitesimal *p/n*, the directions of a “random” rotation should have the same variance as any of the 600 variables involved. But in the high-*p/n* setting the PCA is far from a random rotation, as each PC is limited to the (*n* − 1)-dimensional span of the centered specimens themselves, not the full space of directions among the *p* variables involved. In this example, then, the PC’s are restricted to a 59-dimensional subspace, avoiding all 541 dimensions of precisely zero variance, and yet they must “explain” the total variance of all 600 measurements, which will be about 600. So the PC’s must average variance just over 10, which matches their median in the scree plot. The Marchenko-Pastur distribution is evident in the transposed-ogive form of this scree plot, steeper at both extremes than in its central region. Scree plots of this general form, having nonzero values far above the average variance of the contributory variables throughout their whole length, cannot arise in textbook examples with small *p/n* ratios. The nonzero eigenvalues in this specific simulation range between 5.07 and 17.18, comfortably close to the limits from the theorem’s formula of 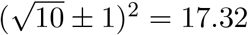 and 4.68.

Then the problem induced by the Marchenko-Pastur phenomenon for large *y* is plain from the panels in the lower row, which scatter the first two and last two PC scores, respectively. Over each panel I have superimposed a circle of radius 2, which one expects to cover most of the variation in any of the scatters at upper left. Instead the spread of the PCA scores is enormously inflated with respect to the underlying variability of these projections, which, recall, were in fact completely arbitrary (because the dimensions of the model that they purport to summarize are actually uncorrelated). The gap between the first and second eigenvalues here is *not* significant by the stepdown test of Bookstein (2014:324), but the visual separation in the upper right panel would nevertheless tempt us to interpret the formula for PC1 anyway; and the segregation of the smallest eigenvalues far away from zero is not part of any of the standard protocols for checking on the realism of PC extractions in the course of GMM analyses or anywhere else in biometry. Hence the meaninglessness of this particular PCA is not accessible to the typical user of PCA software unless that user is fully informed of the consequences of its huge *p/n* ratio. (In fact, for *p/n* ratios near 1, the minimum eigenvalue in the theorem’s distribution formula approaches zero, disabling the separation critique I have touched on here. That infinitesimality leads to a wide range of problems of its own, the central point in my earlier discussion of the MPT, Bookstein 2017.)

**Figure 7.**
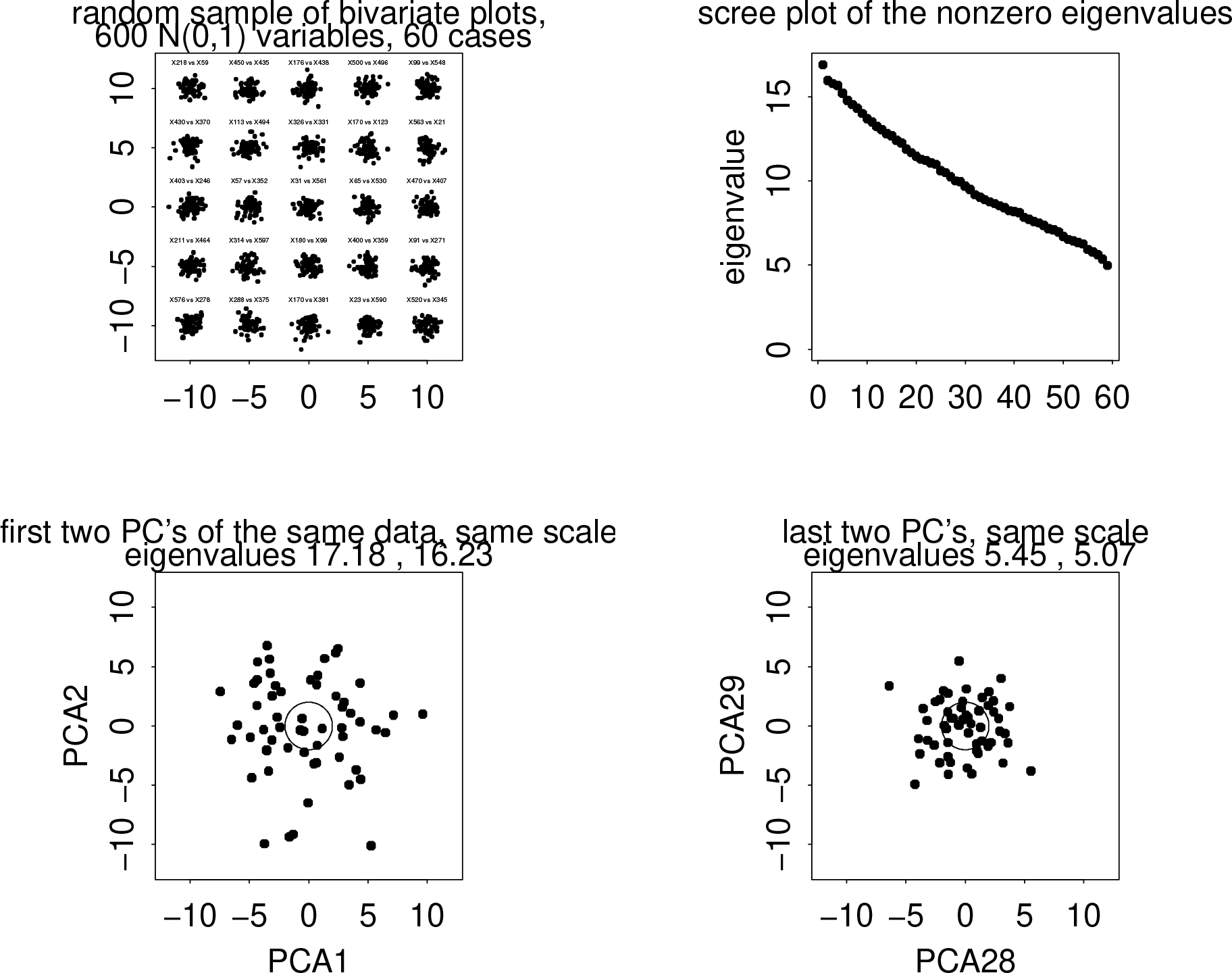
Illustration of the effect of the MPT on an ordinary (not between-group) PCA of too many spherically distributed variables. The simulation here is of 600 independent Gaussians on 60 cases, so *p/n* = 10. (upper left) A random sample of 25 scatterplots of pairs of variables shows homogeneity of these pairwise relationships. The almost unreadable labels simply state the pairing. (upper right) Conventional scree plot of the 59 nonzero eigenvalues. (lower left) Scatter of the two PC’s of greatest variance. (lower right) The same for the last two PC’s. The circle in either panel of the lower row has radius 2; it approximates the covering circle of any of the scatters in the panel at upper left.

Although the MPT is exact only in the limit of very large values of both *p* and *n*, nevertheless I’m invoking it in the small-sample case, indeed the smallest possible sample: just two dimensions being eigenanalyzed. This is what you get if you take *g* = 3 in the notation above. Consider, then, *g* = 3 groups of, say, *m* = 10 specimens each, thus a total sample size of *n* = 30, and a collection of *p* = 300 “shape coordinates” that are independent identically distributed standard Gaussians (mean 0, variance 1). I also need a grouping variable, which I take simply to be the first ten, second ten, and last ten of the thirty cases (except for Figure 14, where this ordering is systematically permuted before being cut into thirds). It will be important in the sequel that the count of these combinations is 30!/(3! · (10!)^3^) = 925166131890 — call it a trillion. ^3^ So step 1 above is done.

For this random subgrouping, compute the *g* 300-vector group means, and assemble them in a “data” matrix, 3 rows by 300 columns. That completes step 2.

Proceed with step 3, the principal component analysis of the three group means as if they were three ordinary single specimens. This is the usual “mean-centered” PCA, meaning that there are actually two dimensions of variation around the grand mean 300-vector. In the general case, this is *g* − 1 dimensions.

Then compute step 4, the imputed scores for each of the original *m* = 30 cases, and step 5, their scatterplots. In those plots, I have numbered the groups, but there is no point yet in numbering the individual cases. (We will need those individual specimen numbers in connection with some of the later figures.)

To ease the application of the MTP’s asymptotic formulas in this setting of only *g* − 1 = 2 dimensions, replace the range 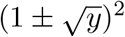 from the MPT by its midrange, which is 1 + *y*, where in this application *y* is huge — 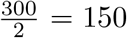 in the example in Figure 1, for instance. The interquartile width of that range is less than 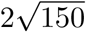, only about one-sixth of its median; I will ignore it. Further, when *y* is this large it is reasonable to omit the “1+” term so as to arrive at a simpler approximation, that ratio *y* = *p/*(*g* − 1) itself. But this was the formula for the PC’s of the theorem’s set of variables of *unit* variance, while our means are of groups of *m*, so we have to multiply the MPT expectation *y* by the variance of those means, which is 1*/m*. So the expectation of our PC variances is now *p/m*(*g* − 1).

Now for one final adjustment. The PCA that I’m applying the MPT to is of the *centered* group means. That reduces the variance of each of the *p* incorporated dimensions by a further factor of (*g* − 1)*/g*. Multiplying, we finally arrive at our expectation of how the bgPCA method should work for this class of simulations: the eigenvalues should be distributed tightly around

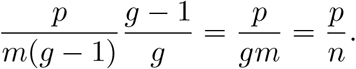

What a simple formula! The expected value of those bgPCA variances is approximately *y* = *p/n*, a new value of *y* that is the ratio of the count of variables to the count of specimens (instead of the count of groups, minus 1). In the simulations of Figure 2, then, the eigenvalues of the PCA’s inside the bgPCA should run about *p/n* = 300/30 = 10. And in the panels of Figures 1 through 4 that is what they do. Note that group size *m* no longer appears in this formula.

Now the origin of the pathology of the bgPCA method in these null models at high *p/n* has become explicit: the unexpected similarity of the scaling of the axes in Figures 1 and 2 to their scaling in the lower left panel of Figure 7. Eigenvalues of the first *g* − 1 principal components of the simulated data will be approximately bounded above by the function 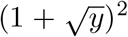 of the design parameter *y* = *p/n* specified by the MPT as modified for this bgPCA setting, while all *g* − 1 of the nonzero eigenvalues of the bgPCA will be approximately equal to this same *y*. The ratio of these is 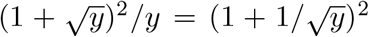, *independent of the group sample size*. For *y* = 10, the value in these simulations, that ratio is 1.316^2^ = 1.73, which closely matches the ratio of 1.718 between the first eigenvalue in Figure 7 and the typical bgPCA eigenvalue of 10 in Figure 2. For the example in Figure 5, with *y* = 40, the match is even closer: 40 for the average bgPCA eigenvalue of group averages, versus an expected maximum eigenvalue of 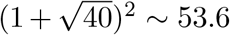 for the underlying data set *in extenso*, for a ratio of 1.34.

This near-equality is unexpected. In ordinary univariate statistics, the variance of a group mean is reduced from the variance of any single observation by a factor of *m*, the group’s sample size. But on our high-*p/n* null model, the variance of the nonzero principal components of the group means is *commensurate* with the variance of the corresponding first *g* − 1 PC’s of the original data, without any division by *m*. Even when *y* is as low as 1.0, this factor 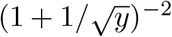 is reduced only to 0.25, more than double the “expected” sample size correction 1*/m* whenever group sizes average 10 or more. The counterintuitive surprise here can be expressed in words instead of algebra this way: the variance within groups (which, keep in mind, remain completely independent of the “measurements” in this class of null models) has dropped drastically from Figure 7 to Figure 1 even as the variance *of* the group centroids has dropped by a much smaller factor — this paradox lies at the core of the pathology of the bgPCA method. As far as we know, the present paper, along with the companion piece by Cardini et al., is the first acknowledgement of this counterintuitive and extremely inconvenient mathematical fact in any printed scientific communication.

The other main pathology highlighted in this paper follows as well from considerations of sample size when it varies across the groups of a single analysis. When all specimens have the same mean, the averages of smaller subsamples have a higher variance than the averages of larger subsamples on every measured (or simulated) variable. The mean vectors of smaller subsamples are thus thrown farther from the grand mean than the mean vectors of the larger samples — the variance of the subsample averages is, after all, proportional to the reciprocal of subgroup sample size. Because bgPCA is an unweighted analysis of those mean vectors, the principal commponents it produces are much likelier to be aligned with the *outlying* subsample averages, which are those arising from the smaller samples. In other words, the smaller groups are far likelier to be the end-members of the between-group principal components. Such a bias of reportage is, of course, inappropriate in any context of ordination, whether evolutionary or not — it confounds the biometric signal with the human difficulty of finding group members as they vary in terms of geography, taphonomy, or geological epoch. As far as I know, this paper is likewise the first acknowledgement of this additional most inconvenient mathematical fact, which is of particular salience to studies of human evolution.

## 3. Factor models

The pathology I am reviewing is not limited to the completely null case, the completely patternless setting, that has characterized the simulations to this point. It also applies to severely distort ordinations in settings where there is a valid dimension or two combined with unstructured noise among a count *p* − 1 or *p* − 2 of residual variables that continues to greatly exceed the sample size *n*. This section considers three such settings, the first two characterized by a single a-priori factor and the third by two factors.

### 3.1. Simulated data with a single factor, possibly with a real example

A data set can have a group structure superimposed over a single biological factor that may, in turn, either be constant within groups or else show some sort of graded phenomenon (such as allometry) there. To say there is only a single factor implies that the residuals from this factor are structureless noise. Then the MPT should apply with its full weight to the multivariate analysis of these residuals. Figure 8 shows how unfortunately persistent this implication is: twelve different simulations correctly detecting a true factor (modeled here as constant within groups and with the factor score for group 2 midway between those for group 1 and group 3). The variation of this true factor is precisely as shown along the horizontal axis of these bgPCA plots. The variance of the fictitious second factor (which is indeed uncorrelated with values of the true factor) is exactly what one would expect from the analysis in Figure 1, since reducing *p* from 300 to 299 will not affect the implications of the MPT.

Figure 9 shows a somewhat different geometrical setting for this same pathology. Now each group has variability on this factor — variability that is correctly modeled here — but also two of the groups have the same average factor score. Nevertheless, as in Figures 1 and 2, bgPCA analysis imputes a wholly fictitious second factor in order to separate groups 2 and 3 while “explaining” the same extent of variance as it did in Figure 8.

**Figure 8.**
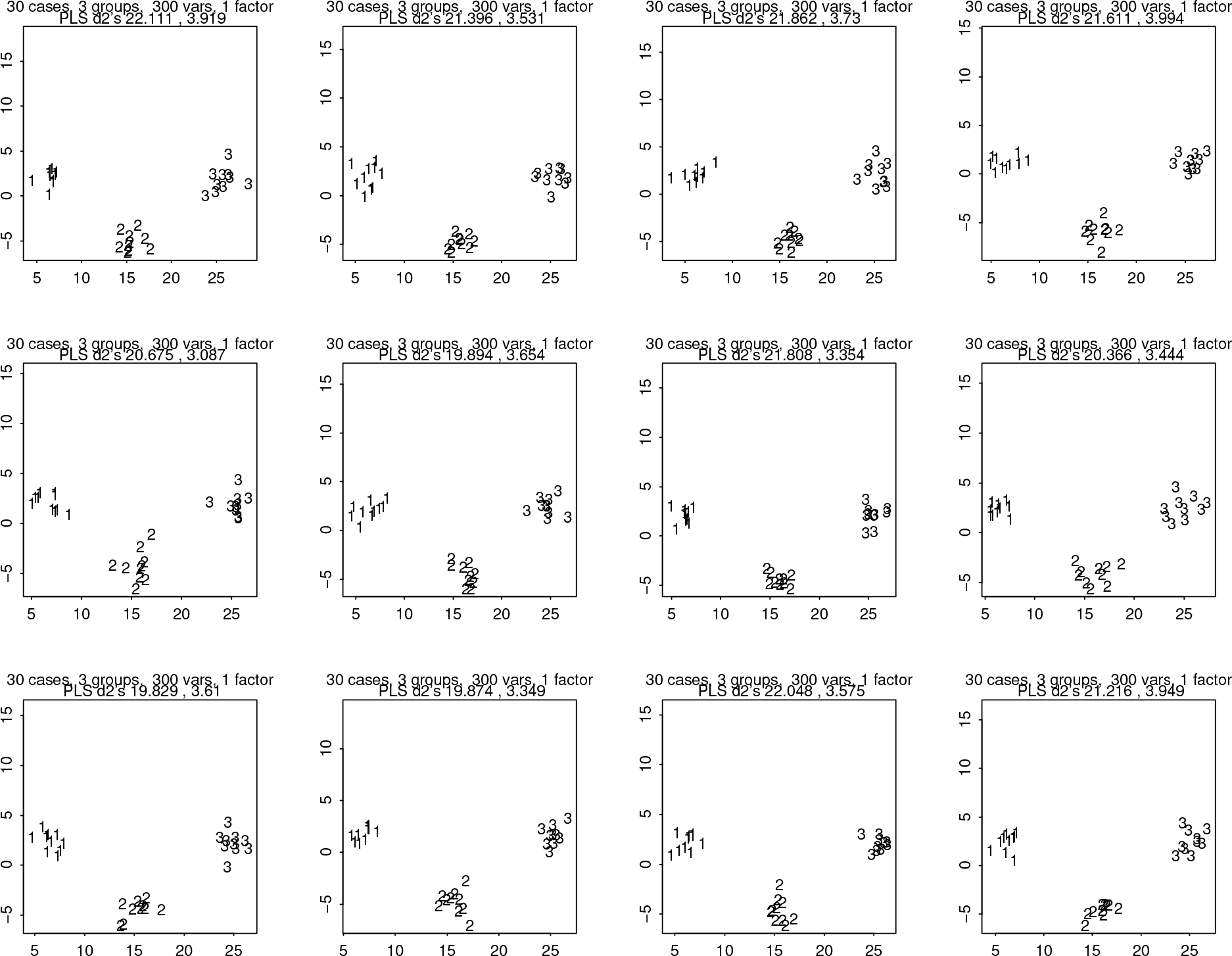
Twelve replicate simulations agreeing on the implications of the bgPCA pathology for a situation with a real (biologically meaningful) factor. The theorem applies nevertheless to the residuals from this factor, pulling out a fictitious second factor with exactly the variance predicted by the MPT, over and over again.

**Figure 9.**
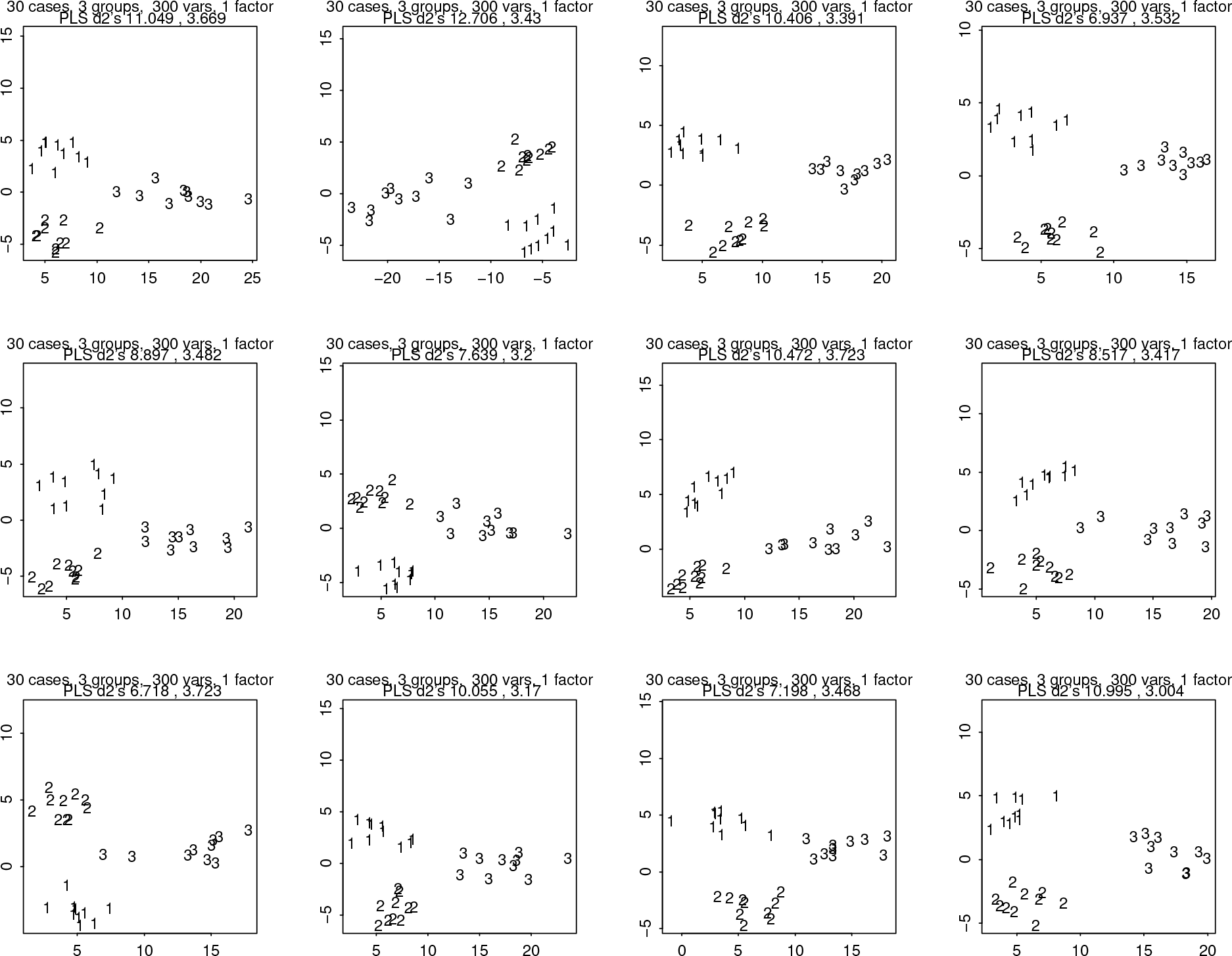
Twelve runs of a modification of the preceding model with two different group means of the true factor score instead of three, along with within-group variance of this score. In every panel the vertical axis is completely fictitious.

The panels of this figure reminded me of a diagram from a paper published in this Journal several years ago: Figure 4 of Mitteroecker and Bookstein (2011). Figure 10 here shows the match after the bgPCA model is tuned to match the *p/n* ratio and the within-group bgPC1 variance of the earlier computation. It thus serves as a realistic example exemplifying this second version of the pathology for an example involving three taxonomic components of the genus *Pan*. In that example, group 3 was separated from groups 1 and 2 but the latter two groups overlapped. The 2011 text near that figure emphasized how much more appropriate an ordination this was than one afforded by CVA, in which groups 1 and 2 separated perfectly. But today the comparison would probably replace the phrase “more appropriate” by a double negative like “less inappropriate.” Back then we did not consider the possibility that the displacement between the means of these two subspecies of *Pan troglodytes* might itself have been an artifact of the bgPCA method just as their perfect separation (shown in another panel of the same original figure) was an artifact of CVA. In this example, *p* was 86, a little less than the total sample size of 104, meaning that the crucial parameter *y* is only 0.83 — the MPT does not yet apply with full force.

**Figure 10.**
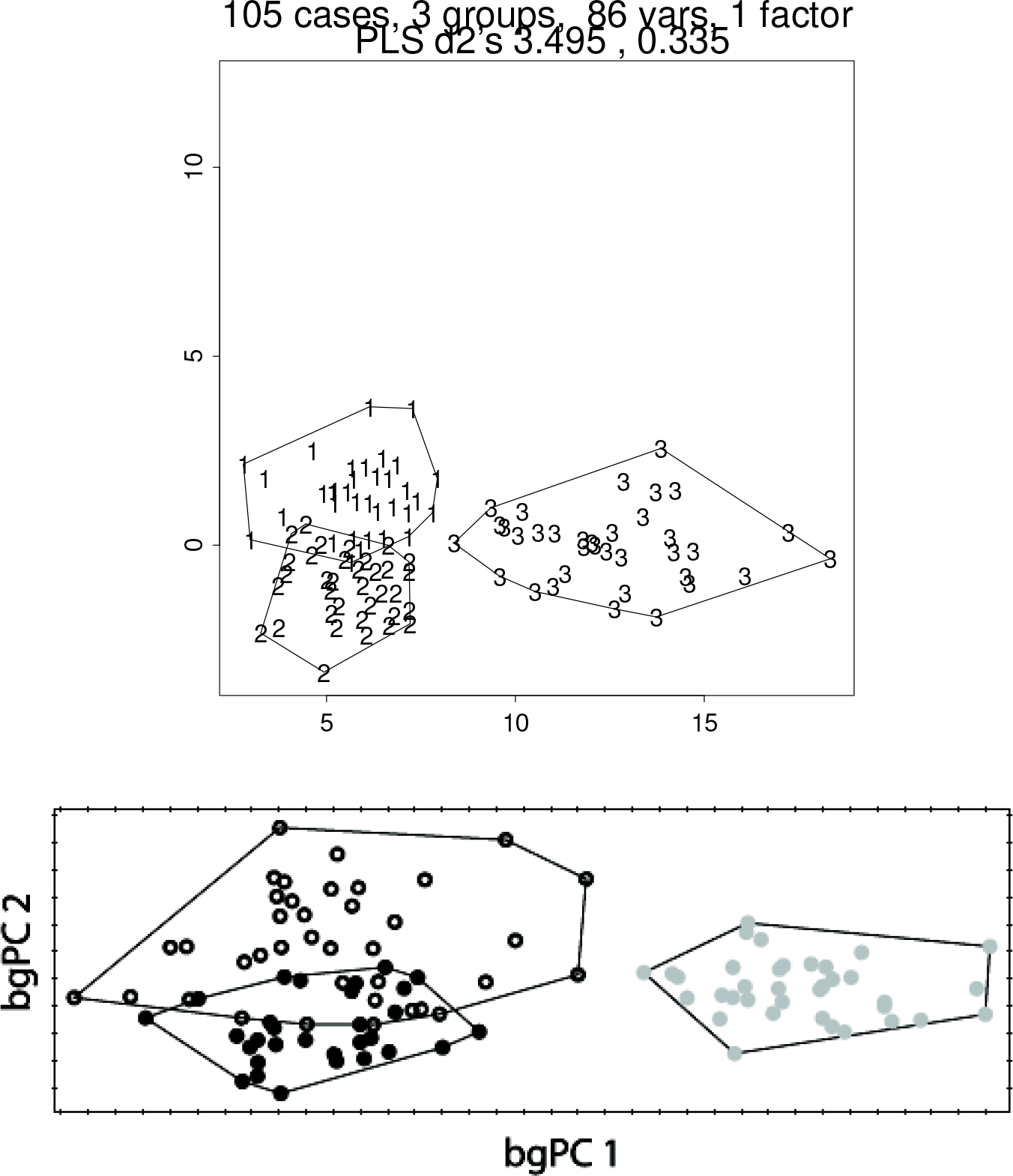
A one-factor model with *p/n* = 0.83 (above) that nicely matches the published two-dimensional analysis (below) of Mitteroecker and Bookstein (2011). The explanatory power of the second decimal quantity in the header is less than that in Figure 1 because the value of *p/n* is only 0.86, much less than the values of 10 or more in my earlier examples. Key to the lower panel: gray circles, *Pan paniscus*; black circles, *Pan troglodytes verus*; open circles, *Pan troglodytes troglodytes. (Permission to be requested.)*

Figure 11 extends this single-factor design to a configuration of *four* groups of ten specimens characterized by factor scores set at four equally spaced values (a reasonable data design for an allometry study in insects, for instance) in a simulation involving 400 variables (so the value of *p/n* continues to be 10). Now bgPCA imagines there to be *two* fictitious factors in addition to the “true” ordination dimension reconstituted as bgPC1. The resulting growth trajectory is parodied as a space cubic over the true (horizontal) axis. The curving cubic trajectory of this factor’s manifestation in morphospace is entirely artifact. Yet the amplitude of its implied polynomial dependence is far from trivial — were it not for our prior knowledge that the *p/n* ratio is so large, we would certainly be tempted to interpret it as a true nonlinearity of allometry. In the rightmost panel of the figure you see a projection of the same regular tetrahedron that was forced by the MPT in Figure 2 (note that the second and third eigenvalues here are nearly equal).

**Figure 11.**
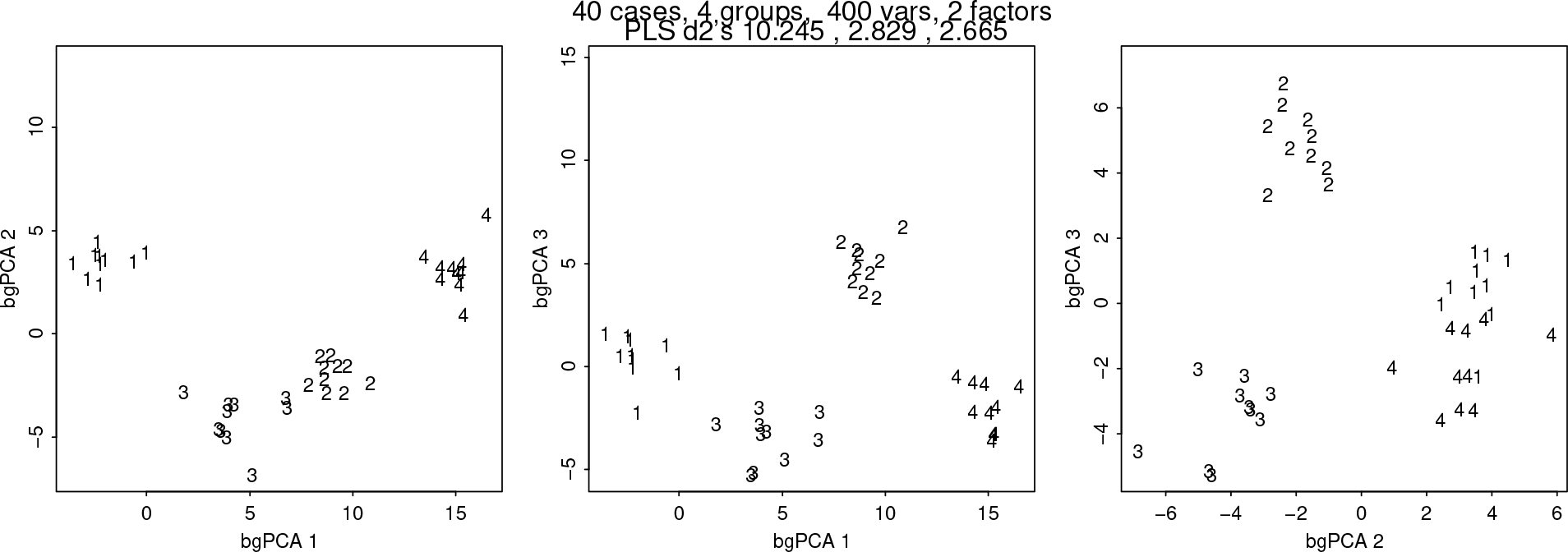
A variant of the single-factor model in Figure 8 results in the completely misleading appearance of a cubic dependence of form on factor score in this 400-dimensional morphospace. Now there are two fictitious factors instead of one.

### 3.2. Simulated data with two factors

Figure 12 extends the preceding example to incorporate two true factors distributed over 400 variables in four groups in a balanced 2×2 design. The value of *y* driving the MPT remains at 10. Again the bgPCA reports three dimensions (one fewer than the count of groups), but instead of the cubic curve in Figure 11 we see a twisted band in morphospace where there should have been a flat rectangle instead. The twist, as you see, is a complete 90^◦^ rotation, from bgPC2 to bgPC3.

In my travels I once encountered a perfectly ordinary object having precisely this form (but in three dimensions instead of 400): a KLM coffee stirrer given to my wife and me on an anniversary trip to Vienna some years ago. See Figure 13.

**Figure 12.**
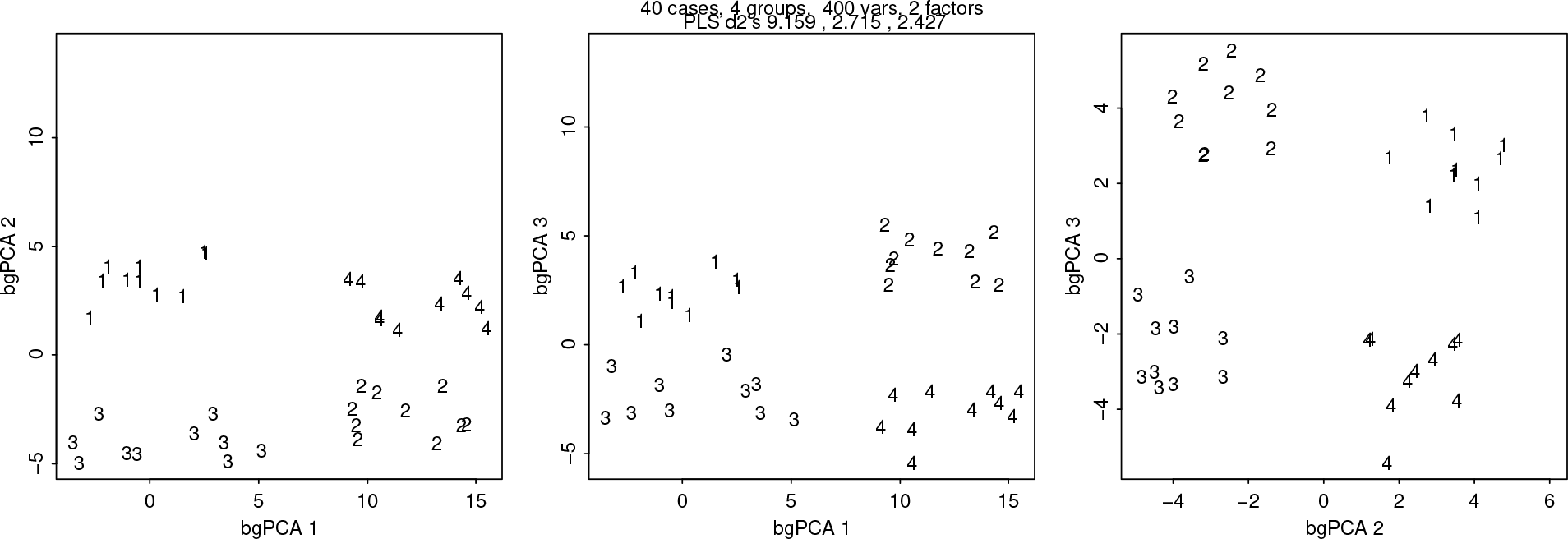
Analogously, a data set with two true factors in a 2 × 2 design is distorted by bgPCA into three dimensions. Note the inversion of the vertical positioning of group 2 vis-á-vis group 4 between the left and central panels — this is the “twist” giving rise to the Möbius strip analogy (next figure).

**Figure 13.**
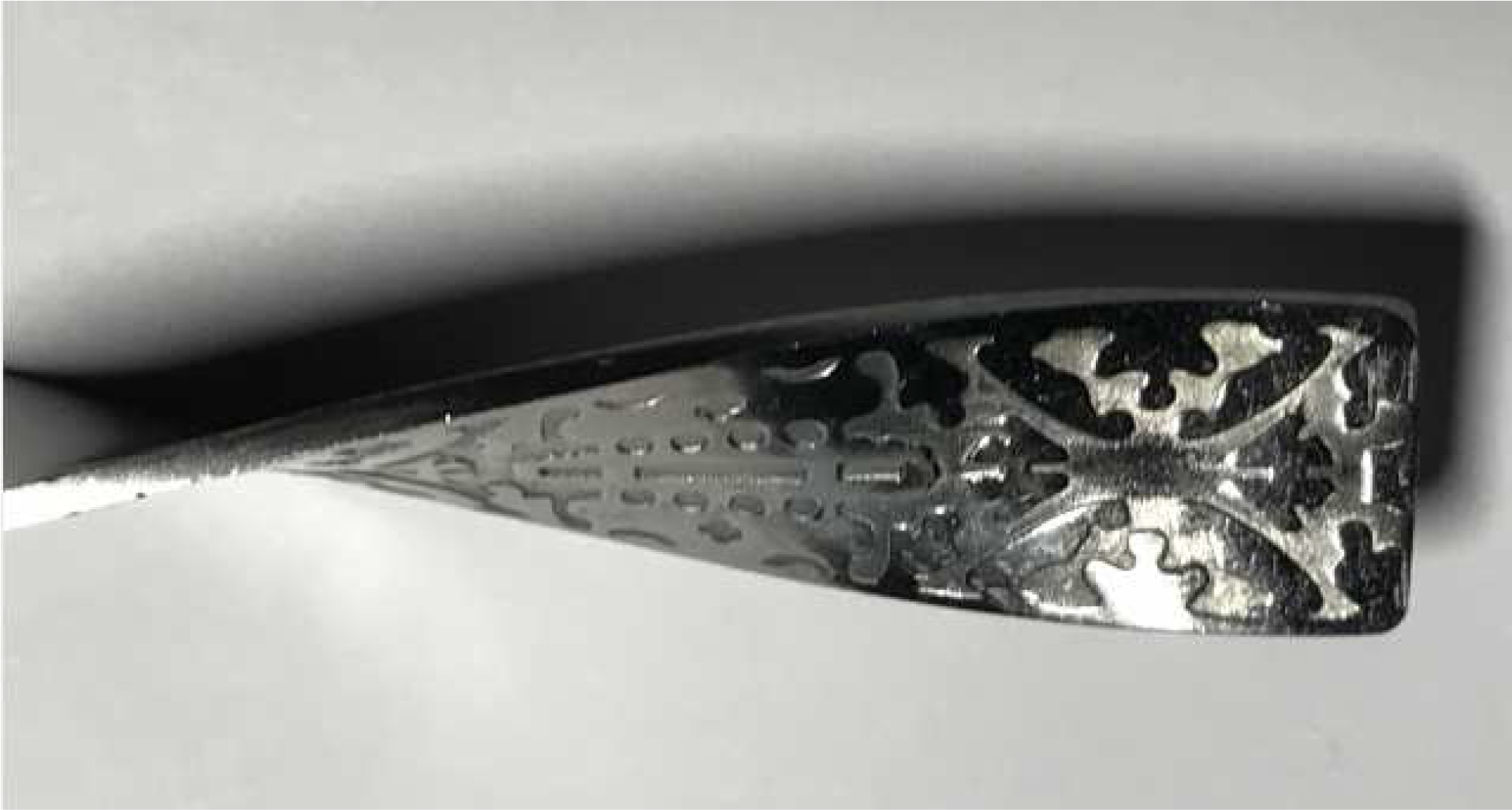
The KLM coffee stirrer, given to me with the compliments of the purser. It is a straightened version of half a Möbius band: the three-dimensional realization of the factor model in Figure 12. The twisting, which here serves a mixed hydrodynamic/aesthetic purpose, is meaningless in the biological context. *(Photo by the author.)*

## 4. Tools for the skeptic

Nothing in the standard textbooks trains biometricians to recognize these pathologies *as* pathologies instead of as publishable empirical pattern analyses. The formal statistical literature of multivariate analysis (e.g. Mardia, Kent, & Bibby, 1979) considers only the context of fixed *p* as *n* tends to infinity, and likewise all the morphometric textbooks prior to 2018 (e.g., Reyment et al., 1984; Bookstein, 1991) deal only with this same conventional large-sample limit. (Reyment’s work elsewhere deals with a great many problems of covariance interpretation, including outliers, but not with this issue of huge counts of variables.) The first mention in the morphometrics literature of PCA’s potentially catastrophic misrepresentations of multivariate reality in the high-*p/n* context is apparently my warning of 2017, and the combination of the present note with its companion piece by Cardini et al. is intended to focus our community’s attention on the single worst case of this pathology, the bgPCA algorithm. The time is appropriate, then, to explicitly review tools that might protect a suitably skeptical quantitative biologist from claiming a separation to be real or a projection of an unclassified specimen to be informative that actually might have arisen from a null data set purely by virtue of its huge variable count. The ordinary machines of significance testing — Wilks’s Lambda, the Hotelling’s Trace statistic, and the like — are inapplicable to the evaluation of multivariate ordinations like these, where misleading claims of separation are built into the very foundations of the ordination machinery. The first two subsections to follow consider *and reject* some standard approaches — permutation tests of group separation and the standard crossvalidation tools of jackknifing and bootstrapping — while the third explores the dependence of bgPCA results on subgroup sizes in more detail. The fourth and fifth subsections briefly sketch two new approaches, one leveraging ideas of factor analysis and the other the technique of shape coordinate deflation, that might show promise.

### 4.1. Permutation distributions: the clusters, or the axes?

Remember that each simulation in Figure 2 is of a *random* subdivision of the *n* = 30 cases into three groups of ten. So, *leaving the 30 points in 300-space unchanged*, produce new random subdivisions, over and over, as in Figure 14. From these unrelated subdivisions you always get the same apparent bgPCA scatter (the same bgPCA eigenvalues, the same separations), albeit variously situated within their square plotting frame. *But the axes of these plots are unrelated* — there is nothing stable about the plane they span.

You can repeat this randomization as many times as you like — the result is always the same. I mean, repeating the subgrouping: there is no reason ever to bother repeating the simulation of those 300 standard Gaussians. You always get a projection of the original 300-dimensional space onto a Euclidean plane for which the projected points coalesce into three grossly separated nearly circular clusters of radius about 1.0 each at unvarying separation. Think of each one of these as the footprint in sand of some equilateral tripod beach stool. A permutation test of the “significance of axis 1,” for instance, would show an enormous significance level for the axis-by-axis ordination actually encountered, in view of the nearly unconstrained tumbling of the bgPCA directions themselves. *And this is certainly the wrong answer*, as for virtually every change of grouping the cosines of the angles the two axes of the new bgPCA makes with the plane of the data-based “bgPC1” and “bgPC2” are nearly as far as they can be from unity. (For the simulation with three groups of ten cases over 300 variables, the median absolute value of these cosines appears to be about 0.17, just a little bit less than 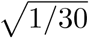.) Then no matter what interpretation a biologist might propound for the axes of the “true” (unpermuted) bgPCA computation, the permutation distribution of the same data set will appear to fail to replicate it, even though the only salient feature (the equilaterality of that triangle of group centers) is replicated almost perfectly.

To circumvent this paradox — that every permutation gives an equally strong signal in an entirely unrelated two-dimensional subspace — would require a specially designed permutation test for uniformity of tumbling as inspected via equilateral separation along with unvarying group concentration per se. No permutation test protocol unaware of the pathologies of the bgPCA method would be capable of assessing the “fit” of this three-group “separation” correctly. Of course there could also be intermediate situations. For instance, a test of the bgPC directions for the analysis in Figure 8 or Figure 9 would need to be subdivided: the direction of bgPC1 should be stable, whereas that of bgPC2 should not be. Analogously, in Figure 11, bgPC1 should be stable, whereas the plane of bgPC2–bgPC3 should be tumbling as wildly as the corresponding plane of bgPC1–bgPC2 in Figure 3 (since in fact they are distributions of almost exactly the same form, random planes tumbling in some 28- or 29-dimensional subspace).

**Figure 14.**
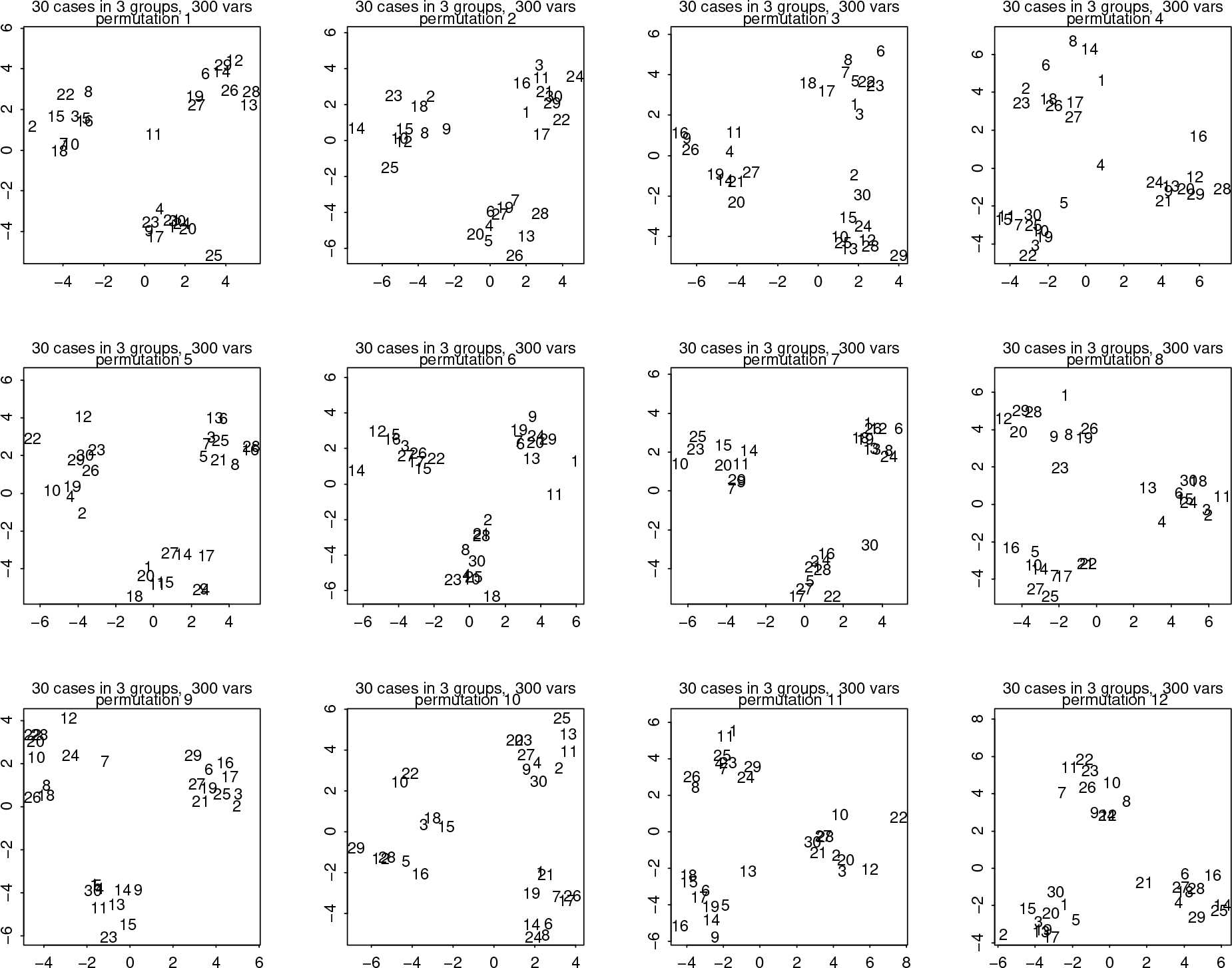
A permutation test of the group code per se for our usual simulation of 30 specimens on 300 uninformative Gaussians leaves the geometry of the usual bgPCA scatterplot unchanged except for rotation and/or reflection, whereas the parameter that has actually been randomized, the plane of the scatter itself, goes unrepresented in the graphic. The random reassignment of specimens to groups is clear in the scrambling of their sequence numbers (1 through 30) in all these replicate plots.

But let me back away a bit from the context of statistical testing to consider the actual algebra of those permutations we are using for purposes of “testing.” Recall that **there were a trillion of these (10,10,10) subgroupings!** So there must be a **trillion** such tripod footprints for the **single** simulation of 30 300-vectors. (Figure 14 showed twelve out of the trillion.) Evidently these are all the same scatterplot, randomly rotated and with the case numbers jumping totally patternlessly between the clusters. In effect there are two ways to manage this permutation test, depending on whether one examines the plane spanned by the two bgPCA’s or instead the pattern of the group centroids, and only when the null model is understood correctly do they yield consistent inferences: the geometry of the centroids is a fixed artifact of the algorithm, whereas the plane of the bgPCA’s is effectively unrestricted in distribution over the appropriate high-dimensional manifold. Such a construction goes completely unmentioned in the standard textbooks of permutation analysis, e.g. Good, 2000.

The argument extends in the obvious way to studies with more than three groups. There are 40!/(40!·(10!)^4^) = 1.96 × 10^20^ (200 million trillion) different permutations of the assignment of 40 cases to four groups of 10 in Figure 5, and they all will result, likewise, in the same three-dimensional scatter, up to orientation and reflection. So this four-group permutation test could concern itself *either* with the three-dimensional subspace spanned by the bgPCA’s jointly *or* with the location of the group centroids within this volume, and again both these approaches, *when interpreted correctly*, affirm the same null distribution.

This is a spectacularly counterintuitive fact about 300-dimensional or 400-dimensional Euclidean geometry, comparable in its import to Gavrilets’s (2004) insights into the dynamic effects of the geometry of edges on high-dimensional surfaces of selection. Indeed it should shock the intuition of any trained biometrician however competent, a shock commensurate with the analogous comment that opens Chapter 3, the chapter on variability of random walks, from Feller’s great undergraduate probability text (it was mine back in 1963 — I still have my copy from then, still bearing its original list price of $9.75). Feller noted that his readers might

> encounter theoretical conclusions which not only are unexpected but actually come as a shock to intuition and common sense. They will reveal that commonly accepted notions concerning chance fluctuations are without foundation and that the implications of the law of large numbers are widely misconstrued.

The situation here is shocking along many of the same lines. The existence of nearly a trillion equivalent tripod-footprint projections, all yielding scientific nonsense, for a wholly signal-free distribution of 30 cases in 300 spherically symmetric Gaussian dimensions is outrageously unexpected, not to mention inconvenient, especially nowadays when it is so easy for the naïve user to produce such megavariate data sets from off-the-shelf imaging software. With respect to this setting the ranges of validity of all the classical exact formulas or approximations to significance levels (Hotelling, Bartlett, Wilks) are simply inaccessible — there are no asymptotics to guide us beyond the MPT as it applies to our familiar canons of multivariate data analysis. One can only conclude that for purposes of inference in this context of GMM with high *p/n* ratios (in fact, I would argue, in *any* multivariate context) *permutation tests should be used only to accept hypotheses, not to reject them*.^4^

### 4.2. Crossvalidation

The preceding argument, while phrased in a way specific to permutation testing, serves more generally as a critique of the standard logic of crossvalidation by a variety of resampling techniques (regarding which in general see Efron 1987). Before we can apply it to “test” findings in this domain of high-*p/n* morphometrics, we need to decide what specific aspect of the arithmetic it is our concern to validate. Is it the exploitation of the classification arithmetic that is our central purpose, or the furthering of our understanding of the ordination by its conversion into an explanatory scheme of empirically stable dimensions that will eventually be interpreted as factors? In other words, is the scientific deliverable of the analysis that led to figures like Figure 1 the production of the scores or instead the production of the scientific explanations, whatever they might be, suggested by the alignment of the bgPCA axes? (The problem is not specific to the bgPCA context, but arises whenever a biplot or the singular-value decomposition driving it is invoked in the course of any empirical data analysis.) The literature of bgPCA is silent on this question. Yendle and MacFie (1989:589), for instance, set their discussion of what they call DPCA as a tool for performing “a supervised pattern recognition,” the production of “factors that describe the between-group variation most effectively.” It is left unclear whether that most effective pattern description deals with the scores or instead with the axes.

In my view, our concern as users of GMM must be with the axes, not the scores. To note that group 1’s scores in Figure 1 average 3.9 on axis 1 would not be a helpful observation. This is because we are working in a context of megavariate analysis, where there is no theory governing the scientific meaning of the specific selection of variables — there is no answer to the question, “3.9 on what scale?” No, the individual and average scores, arising as they do from a theory-free list of shape coordinates, have no numerical meaning in and of themselves. As I have argued in some detail in Bookstein (2019), intellectual progress under such circumstances mandates an analytic rhetoric taking the form of some sort of *explanation*. The foundational literature of PLS was occasionally quite explicit on the topic, under the rubric of the “reality of latent variables” (see, e.g., Bookstein, 1982). The PLS analyst’s attention focuses on the interpretation of the axes in Figure 1 not as principal components of anything (even though that is how they are usually computed) but as “latent variables,” quantities that should be interpreted as if someday they would be capable of explicit measurement.

Here in 2019 the battle between these two thrusts in contention, exploitation of scores versus interpretation of axes, is over, thanks to the enormously successful new discipline of *machine learning* (see, e.g., Hastie et al., 2009). The huge range of current silicon-intensive approaches — kernel smoothing, support vectors, neural nets, and many others — has jointly rendered linear classification methods wholly obsolete as a component of competent contemporary empirical natural science. (In particular, canonical variates analysis and its special case, linear discriminant analysis, are no longer worth teaching except as historical curiosities.) Then the claims that need crossvalidation are those related to interpretation of the *axes* in the figures here, not to quantifications of the group assignments entailed, and the versions of crossvalidation that apply would be search algorithms through data sets near the instant data set in a variety of respects, not any single formula.

I think the first example of such an application to the present analytic setting (this one a bootstrap, as the context was not one of subgroup analysis) may have been one of mine of three decades ago, Sampson et al. 1989. Briefly, jackknifing is the recomputation of analyses *n* times, leaving out each one of the *n* specimens in turn, while bootstrapping is the indefinite reanalysis of subsamples of the same total count *n* made up by resampling *n* items from the original data set “with replacement,” meaning that individual specimens can appear more than once. The purpose of either is the estimation of the standard error of a parameter from the true model: for further background consult Efron 1987. In either context, while typically a permutation test is aimed at showing a range of reanalyses that *fail* to agree with the “true” computation, a crossvalidation approach is intended to show a range of reanalyses, or even a complete census of them, that instead *agree* — that result in almost the same pattern claim.

Thus to apply any crossvalidation technique to an investigation to which the null hypothesis of sphericity is relevant we must decide on a criterion for when two or more analyses are “almost the same.” When equality of eigenvalues is a possibility, two eigenanalyses can be said to “match” if either set of eigenvectors can be rotated (with or without reflection) onto the other. (This was actually the application for which the psychologists invented Procrustes analysis in the first place, as in discussions of the invariance of the pentadimensionality of personality scales. See Hurley and Cattell, 1962.) When modified by invocation of the Procrustes procedure in this charming non-GMM context, the fictitious production of an equilateral structure of cluster centers in Figures 1 through 6 proves most unfortunately robust. The pathology explicit in Figure 2, in other words, would be misinterpreted as a confirmation of the original ordination.

Figure 15 is a typical example of jackknifing on our megavariate null model. The thirty frames here arise from thirty separate analyses of the same simulated data set, each one omitting one of the original thirty specimens of the usual 300-vectors of independent Gaussians but then, in order to control the spinning so apparent in Figure 2, rotating each of the resulting configurations to the original bgPCA of all thirty specimens using the rotational part only of the standard Procrustes algorithm (e.g., no centering, no rescaling, thus only one degree of freedom exploited instead of the usual four). You see how stable the fiction is against this type of challenge. The jackknife formula for the standard *error* of a parametric summary of this type declares it to be the standard *deviation* of the parameters over the jackknifed resampling. In terms of the axes of the diagram these derived standard errors are 0.124, 0.040, 0.057, 0.164, 0.071, 0.184. So these cluster locations, and presumably the corresponding organismal interpretations of whatever contrasts the underlying plane of dimensions invokes, would be inferred to be stable against accidents of specimen sampling. Ironically, that inference is correct — the computed geometry of the group centers *is* that of an equilateral triangle, tetrahedron, etc. — but this is a property of the algorithm per se rather than an empirical finding saying anything about actual biological data.

**Figure 15.**
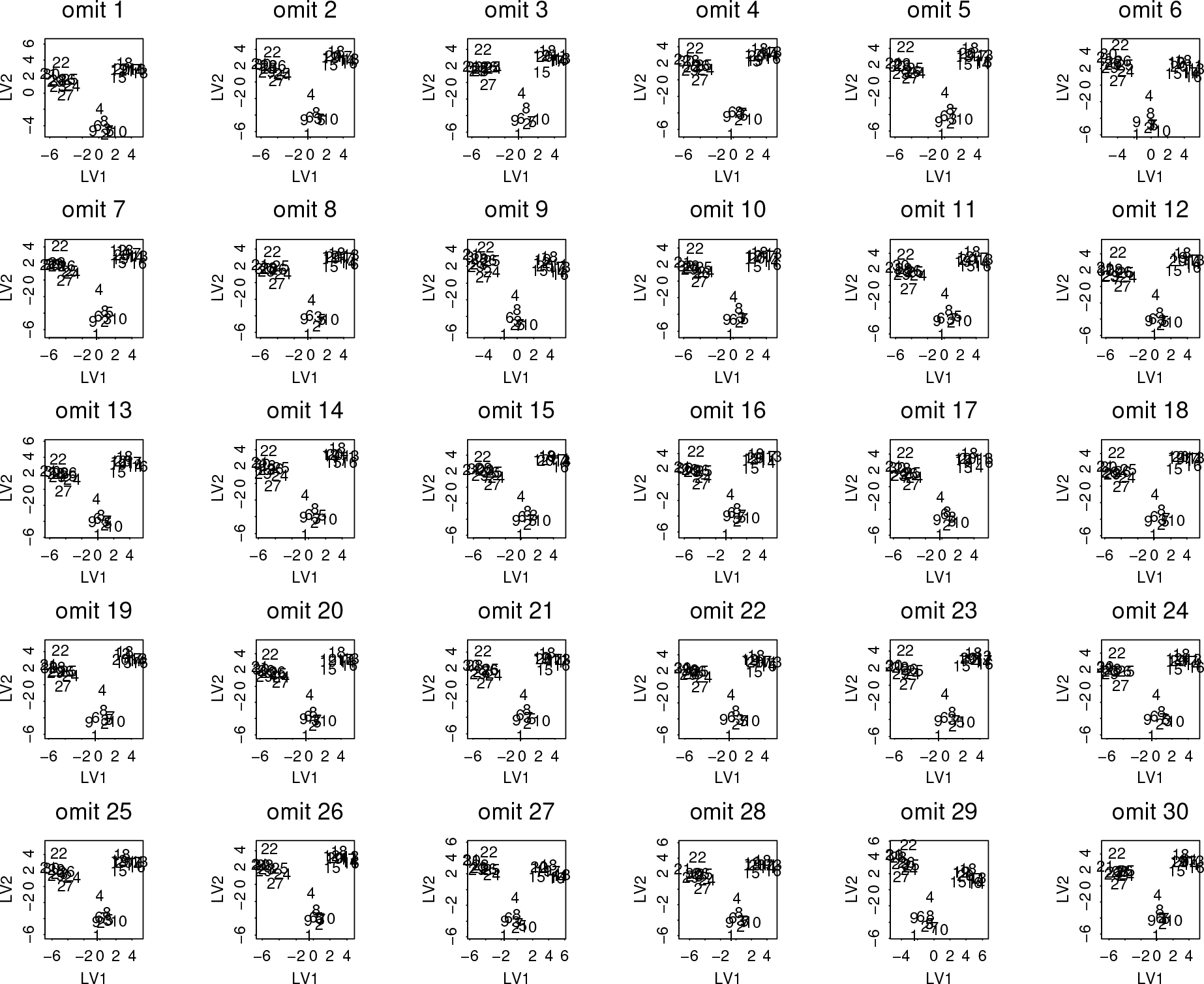
Applying the classical “jackknife” technique to the scenario in Figure 1 or Figure 2. Each panel is the analysis of one of the 29-specimen subsets from the usual simulation of totally uniformative Gaussian noise on 300 measurements over 30 specimens. Printed numbers are specimen numbers matching from panel to panel.

Applying an alternative standard resampling approach, bootstrapping, to bgPCA analyses in this null setting proves interesting. The example in Figure 16 concerns a scheme of 60 specimens over 600 Gaussian variables completely uninformative about the grouping of the data into three groups of 20. The standard bgPCA (left panel) shows precisely the usual fictitious group separations expected when *p/n* = 10.

In the center panel is an inappropriate analysis, a standard bootstrap (resampling specimens with replacement): inappropriate because the bootstrapped version has intrinsically different symmetries. The clusters that had been circular are now strongly stretched in a radial direction, and the stretching seems entirely due to the specimens that have appeared more than once in the resampling (here, printed with jitter so they appear to be in heavier type). In a PCA-like context like this one, sampling with replacement is inappropriate, as any specimen appearing twice in the resample has twice the weight of the nonduplicated specimens in the analysis, three times the weight if it is tripled, four times if it is quadrupled. All such reweightings diminish the variance reduction of the group average and thus increase the leverage the specimen in question has upon the variance of the bgPC’s (as represented in these scatters by distance from the centroid, hence the elongation of the clusters that were circular in earlier figures). This accidental entanglement of the logic of bgPCA with the logic of bootstrapping does not affect the fictitiousness of the inferences here even as it proceeds to destroy one of the symmetries of the MPT-constrained plot in the left panel.

In the right-hand panel of Figure 16 is a more appropriate resampling method that suppresses the overweighting of those repeated specimens: a decimation analysis that randomly deletes a substantial fraction of the sample. I’ve realized it here simply by deleting duplicates from the bootstrapped samples; for groups of 20, this is very nearly the same maneuver as analysis using half the original sample. Unexpectedly, up to rotation and reflection this does not seem to blur the quantitative structure of the resulting geometry of group centers at all — in a high-*p/n* application like this, the effect on apparent precision of proportionately reducing sample size on an unchanging variable count *p* is less than the “improvement” dictated by a corresponding increase in the *p/n* ratio. This stochastic invariance leads to even more strongly misleading equilateral expected circular separations spinning in exactly the same meaningless way. Notice that in both Figure 15 and Figure 16 the role of the crossvalidation, whether jackknifed or bootstrapped, is to support the hypothesis embodied in the simulation model, not to reject it.

**Figure 16.**
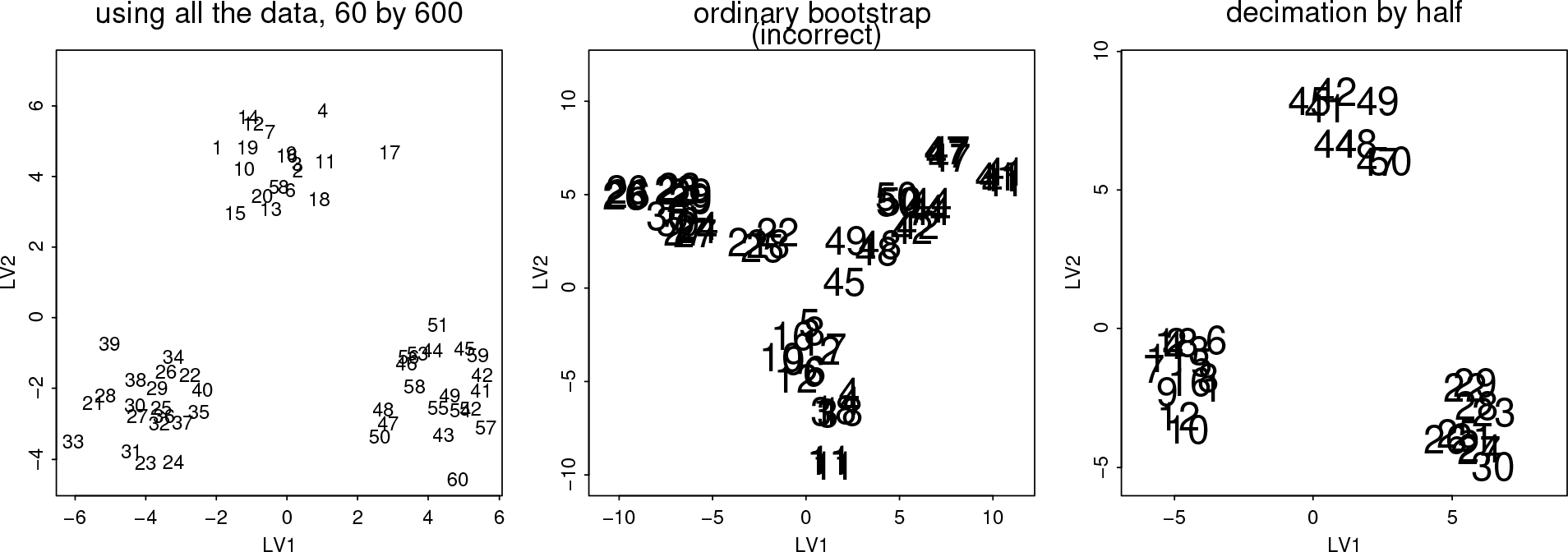
Bootstrapping destroys one of the symmetries of a high-*p/n* null bgPCA, but sample decimation does not. (left panel) bpPCA of a simulated data set of 60 specimens on 600 variables. (middle panel) bpPCA of a bootstrapped resampling, with duplicates randomly jittered to increase visual weight. Notice the change of scale from the first panel. Duplicates in the course of resampling are printed with jitter. The inappropriate weighting assigned to the duplicated specimens is clear here. (right panel) Replacement of the bootstrap by a simple decimation (here, by half) yields not the customary reduction of precision but instead a modest intensification (note the scale change of these axes) of the original cluster center separations at left owing to the concomitant doubling in the value of *p/n* from 10 to 20.

### 4.3. A pathology arising from inconstancy of group specimen counts

When Section 2 introduced the formulas that account for the simulations of this paper, it noted that “for simplicity” I would set all groups to the same sample size. It is convenient here to return to that assumption, because the effect of varying group size is identical to the effect of varying individual specimen weight in the bootstrapping approach that was just highlighted in Figure 16. Consider one such example in which sample size varies realistically: a total of 130 specimens of which 10 are from group 1, 30 from group 2, and 90 from group 3, “measured” on 650 standard Gaussians independent of group, so that *y* = *p/n* = 5. As you can see from the usual bgPCA scatterplot, Figure 17 (left), the smaller the group, the farther its projected bgPCA average score from the grand mean, just as in the central panel of Figure 16. The most obvious symmetry of the bgPCA analysis for a null model has been broken merely by accident of sampling design.

One might attempt to fix this problem by resampling each of the larger groups down to the sample size of the smallest, much as was done (approximately) in the right-hand panel of Figure 16. (The resampling would need to be done repeatedly if there is some possibility of heterogeneity within the larger groups.) But such a jury-rigged adjustment to bgPCA fails in the presence of the even more severe inequality of group sizes that are typical of studies combining extant and extinct species, or *Homo* with other extant primate genera, or human populations sorted geographically from samples of burials or from museum collections. At the right in Figure 17 is the bgPCA analysis of a simulation of four imbalanced groups of sizes 40, 10, 7, and 3 on 300 wholly uninformative “measurements” (so the *p/n* ratio is still 5). In paleoanthropology, for instance, such a range of group sizes is typical for studies that combine samples of *H. sapiens* with other species of *Homo* or with genera of extinct anthropoids. In this setting, under the usual null model the smaller groups will typically dominate every reported between-group principal component, in order of group size — here the (fictitious) contrast of group 1 with the smallest group, group 4, drives bgPC1, while that of group 1 with the next smallest group, group 3, drives bgPC2. It would be fallacious to argue that group 4 embodies some sort of apomorphy based on its position here when that position is explicitly a function of our failure to locate more than three specimens of that group beforehand. Surely no such analysis deserves to be described as an ordination at all. Compare the count and the position of the *H. erectus* specimens in bgPCA scatterplot Figure 4c of Détroit et al. 2019.

Recall the formula from Section 2 that the factor by which bgPCA inflates the variance of a bgPCA component over its expected attenuated value is roughly 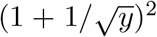. For *y* = 1, this is 4.0. Then for there to be any degree of validity in subsequent inferences, the size of the groups whose positions are the topic of inference in bgPCA scatters from such data designs should be more than double this correction factor, or about 10, whenever the *p/n* ratio is 1.0. For higher *p/n* this stricture will be even more severe. For instance, for group 4 to lose its misleading dominance in the right-hand panel of Figure 17, the eigenvalue of bgPC1 would need to drop below that of bgPC2 here, which approximates the design parameter *y* = *p/n* = 10 of the simulation. To shift the mean of group 4 so that its contribution to the variance in this direction no longer swamps that basic effect of high *p/n*, it would have to move left to a position of roughly 4 on the abscissa instead of 10. This requires an increase of sample size by a factor of about 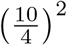. But somewhere along this path of increase in sampling frequency it will cease to be the smallest group, and axis bgPC1 will jump to some other alignment instead.

**Figure 17.**
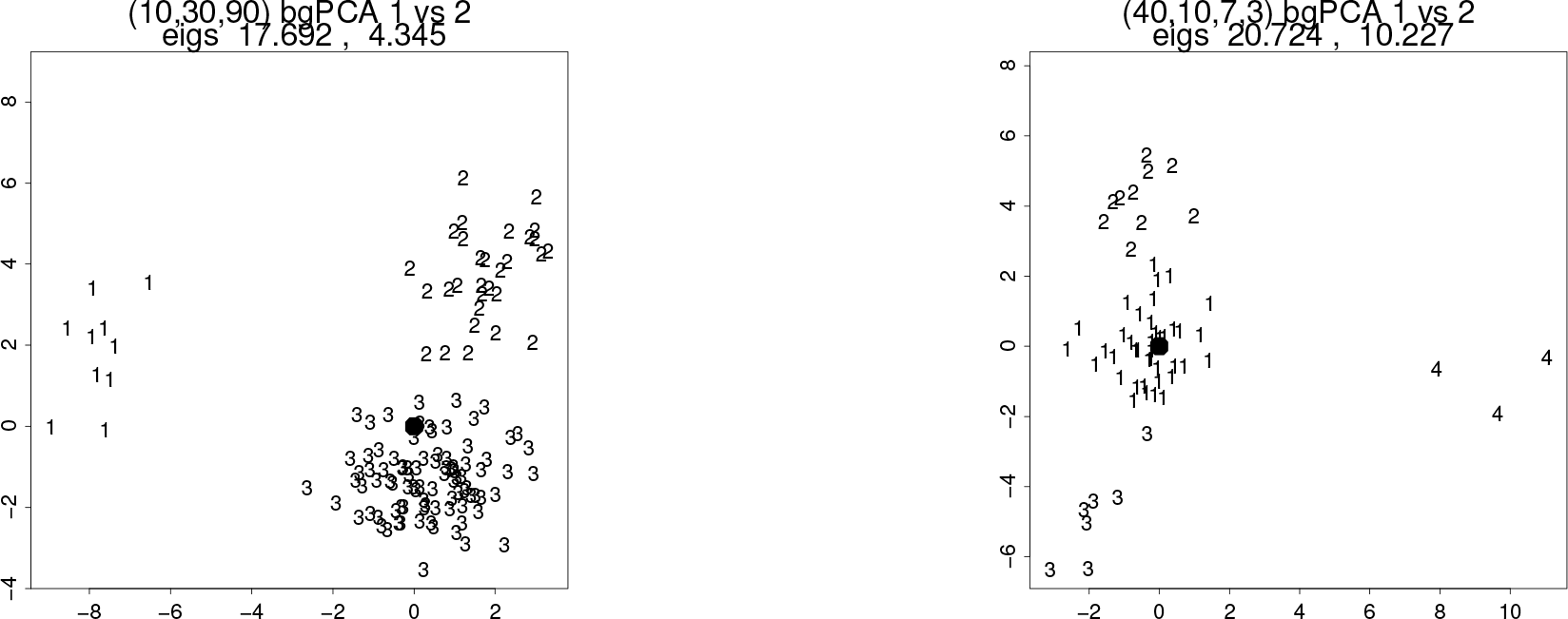
The effect of variations in group size is exactly analogous to the effect of multiple appearance in a bootstrapped analysis. (left) A bgPCA of 650 standard Gaussians for a sample of 130 specimens in three arbitrary groups of sizes 10, 30, and 90 throws each group average to a distance from the grand mean (the large filled disk) proportional to the inverse square root of its group size, thereby breaking one diagnostic symmetry of the bgPCA critique here. (right) Even more extreme disproportions lead to further pathologies of a bgPCA report, here, from samples of size 40, 10, 7, and 3, respectively. See text.

Figure 18 explores such an evolution in more detail by extending the preceding simulation in order to systematically vary the ordering of subgroup sizes. Here the three largest subgroup sizes from the right-hand panel of Figure 17 are preserved, but the count of group 4 is modified in five steps, one that actually reduces it from 3 specimens to 2 and four others that increase it to 5, 10, 15, or 30 specimens, respectively. Each simulation is of the appropriate total count of specimens (from 59 to 87) on five times as many totally uninformative standard Gaussian measurements. At size 2 (upper left) the alignment of bgPC1 with group 4 is even more unequivocal than in Figure 17, while bgPC2 continues to be closely aligned with the *second*-smallest group, which here is group 3. Note also that the distance of group 4 from the grand mean is roughly twice that of group 3, corresponding to the square-root of the ratio of subgroup counts. At upper center, the size of group 4 is nearly the same as that of group 3, and so the distances of the subgroups from the center of the bgPC1–bgPC2 scatter are nearly equal, and likewise their leverage on the first bgPCA axis pair. In the simulation at upper right, the size of subgroup 4 has been set to 10, which is no longer the smallest, and so bgPC1 has jumped to an alignment with group 3 instead. But as now all of the subgroups except for group 1 have roughly the same specimen count, the bgPCA becomes the projection of a tetrahedron onto the face opposite group 1, with the largest group at the center (actually, it’s an end-member on bgPC3, not shown) and all three of the other groups as vertices of the projection. At lower left the count of group 4 is now 15, breaking the symmetry with group 2, so the bgPC1–bgPC2 axes are now a rotation of the displacements from group 1 to group 2 and to group 3 at roughly 90°; in comparison with the panel above, the role of group 4 has been supplanted by that of group 2. Finally, at lower center is a simulation with group 4 set to 30 specimens. Groups 3 and 2 continue to determine bgPC1 and bgPC2, respectively, and groups 1 and 4, the two largest, project on top of one another here because their contrast determines precisely the direction of bgPC3, which is scattered against bgPC1 in the lower right panel.

Hence the bgPCA scatterplot of a high-*p/n* simulation of the null model over groups that differ widely in sample count can be predicted almost exactly from those group counts alone, independent of every bit of information in the simulated “measurements” except for the value of the ratio *y* = *p/n*. No such scatterplot could possibly contribute to any valid inference unless and until this null model, or, more likely, its combination with within-group factors, is unequivocally rejected.

**Figure 18.**
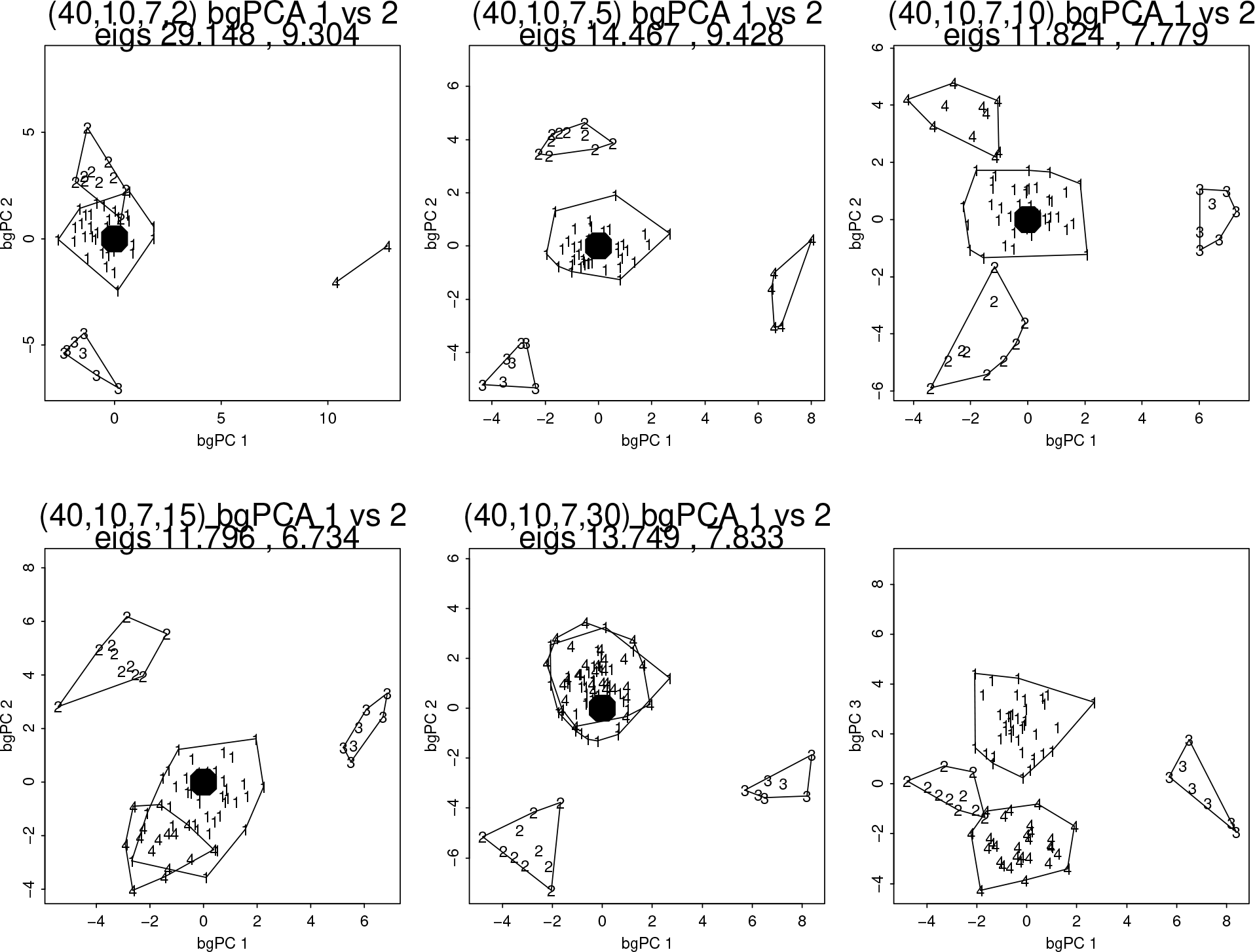
The pathological effect of subgroup sample size variability as demonstrated by a systematic manipulation of one group size out of the four. Large filled dot: grand mean of the bgPC1–bgPC2 scatter. Every ordination in this set of panels is fictitious, in particular the apparent separations of the clusters in every panel. See text.

### 4.4. Some aspects of truth persist in spite of the pathologies

When applied to a high-dimensional null distribution, the fictitious group clusters that bgPCA creates resemble Gaussian disks or balls. Scatters in which the group clusters are elongated, then, are unlikely to be expressing solely the bgPCA fiction; the elongation would instead correspond to factors in some true model other than the model of purely spherical Gaussian variation of very high dimension. Figure 10 already showed an example where these Gaussians were shifted by a single factor whose scores varied both within and between groups. In Figure 19 we explore two variants of this approach, in both of which there continues to be at least one true within-group factor, but now without any group differences in distribution. *When the signal from the true factor(s) is strong enough*, the bgPCA algorithm produces scatters consistent with the existence of such a factor or factors and with the overlap of true scores by group. Unfortunately it also produces fictitious factors of group separation as well.

The upper row of the figure shows the bgPCA analyses for two single-factor simulations that modify our standard *p* = 300, *n* = 30 simulation by a single additional factor with loadings Gaussian of variance 0.25 around 0.0. Component bgPC1 of the simulation at upper left correlates 0.95 with the actual simulated factor score here, and (correctly) fails to separate the groups, while as usual there is a completely fictional bgPC2 suggesting a separation. Similarly, the score on bgPC1 from the simulation at upper center correlates 0.83 with the actual simulated factor score, and likewise correctly fails to separate the groups, whereas once again component bgPC2 is wholly fictional.

The lower row of the figure reports a single simulation of four groups, now in a *p* = 400, *n* = 40 design driven by two factors, the first with loadings Gaussian with variance 1.0 around zero (i.e., four times as powerful as it was in the simulations of the upper row), the second with the same variance as the single factor simulated in the upper row. The first bgPCA component again correctly reflects the lack of separation of the groups on this pair of dimensions, correlating 0.96 with the simulated factor score. The corresponding first eigenvalue is too large to have arisen from the null distribution discussed in Section 2.

But the bgPC2–bgPC3 scatter (lower right panel in the figure) once again succumbs to the bgPCA algorithm’s relentless confabulation of group separations. The first of these directions seems roughly aligned with a real factor, but not the second; the separation concocted by the bgPCA algorithm in this panel confounds the two in a fictitious ordination that rotates the direction of bgPC2 away from the actual factor direction as modeled. Because this bgPC2 evidently was computed to accommodate the (fictitious) separation of group 2 from group 4, it correlates only 0.73 with the actual modeled factor score, thereby invalidating the device of projecting an unknown specimen onto the *left*-hand panel, the bgPC1– bgPC2 panel, as any sort of valid inference of affinity. Inasmuch as the lower left panel of Figure 19 generally resembles Extended Data Figure 6 of Chen et al. (2019) except for a squaring of the plot, it would have been appropriate for that paper to have displayed the scatters of all pair of bgPC’s, not just the first pair, in order that readers might check for this possible hidden confound induced by bgPCA’s tendency to generate fictitious separations.

**Figure 19.**
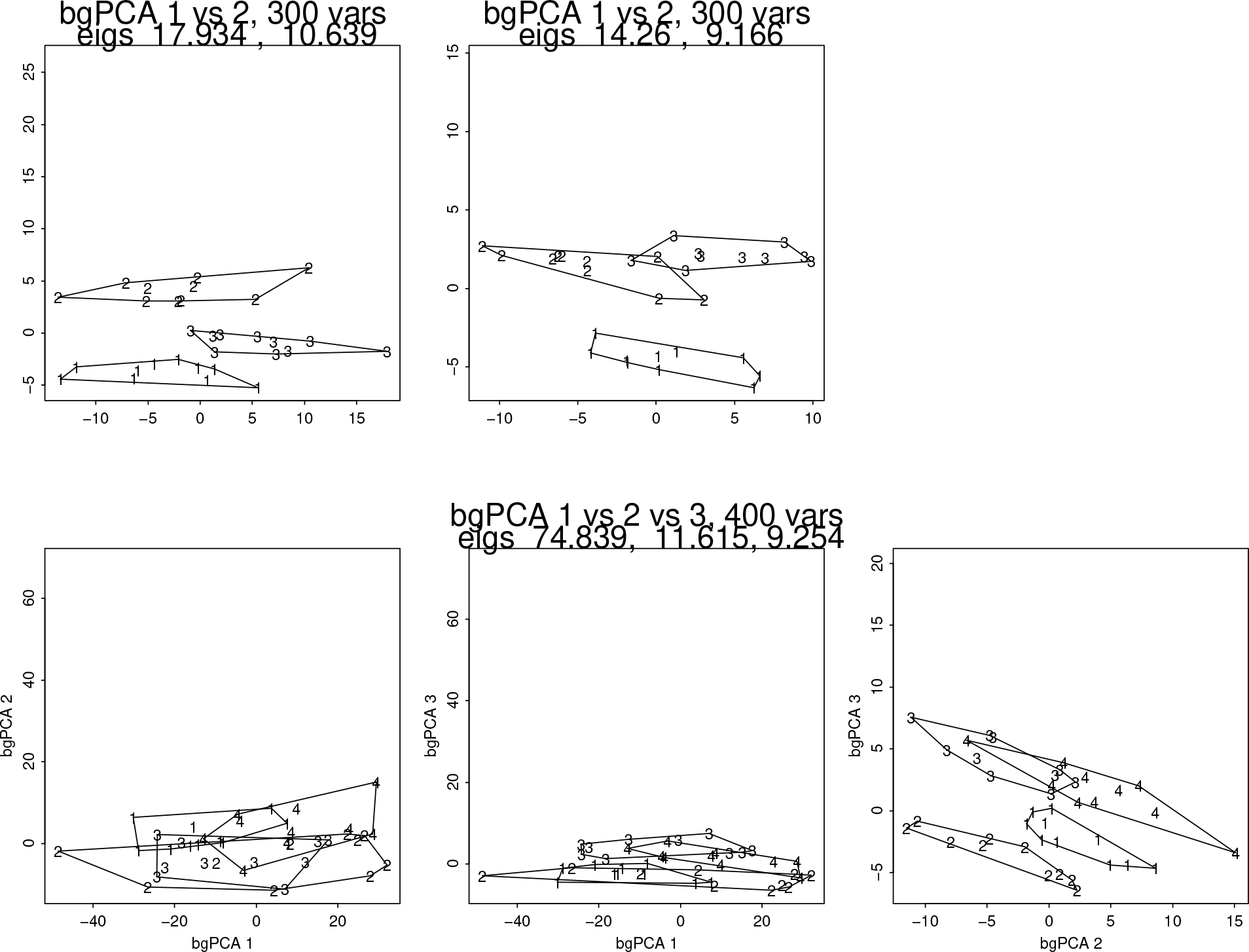
True factor models can coexist with the pathological separations implied by a high-*p/n* bgPCA. (upper row) Two different bgPCA simulations of three 10-specimen groups on 300 Gaussian variables that also reflect a single underlying factor, everything independent of group. In either, bgPCA tries but fails to pull apart the groups on bgPC1, the valid ordination dimension, but succeeds with the fictitious bgPC2. (lower row) The same for a single simulation of four ten-specimen groups over 400 variables, now with two valid factors, likewise everything independent of group. Now bgPC1 and (bgPC2–0.5×bgPC3) are both valid factors, while (bgPC3+0.5×bgPC2) is a 2D projection of the usual 3D fiction. The equality of the model’s group means notwithstanding, the bgPC2–bgPC3 panel separates these (identically distributed) groups distressingly well.

A comparison of these panels to those in Figure 10 suggests the following two simple, if crude, generalizations: bgPCA clusters aligned with apparent within-group factors, and thereby less likely to show group separations, are more reliable than those orthogonal to such within-group dimensions; and bgPCA dimensions showing high within-group variance are likelier to be aligned with true within-group factors. This is, of course, exactly the opposite of the reason for adopting approaches like bgPCA in the first place — the suppression of “bias” deriving from that within-group factor structure — and likewise disrespects the logic driving classical approaches such as Mahalanobis distance, the formula for which downweights the dimensions of largest within-group variance instead of paying special attention to them in this way. One might say, again risking overgeneralization, that analyses of group average differences that fail to attend to within-group variability are not likely to sustain sound biological inferences in high-*p/n* settings such as GMM. It would follow, likewise, that the tests for stability of any “real” factors, like those that show directional enhancement of variance in Figure 19, would need to be separate from the tests for stability of the dimensions in which the clusters by group appear not to be stretched (I have already mentioned this ramification in the comment preceding Figure 14). This further supports the claim in footnote 4 that statistical significance testing is appropriate in high-*p/n* biological sciences only in the vicinity of a true model, that is, after you know nearly everything about how your hypothesis relates to your data except possibly the values of a few decimal parameters.

### 4.5. A potential defensive resource: integration analysis

In addition to low-dimensional factor models, there are other promising approaches that exploit practical experience to constrain the nature of the GMM variability being summarized. Some replace the null model of spherical Gaussian symmetries by an alternative in which the structure of empirical shape variation, whether or not the groups are real, is represented by a range of patterns at one or more distinct spatial scales. There is a taxonomy of these scaled approaches in Bookstein (2019).

One of these, the approach via loglinear rescaling of the BE–PwV (bending energy – partial warp variance) plot, is illustrated here in Figure 20. The figure is based on a single simulation of 30 specimens in the usual 10-10-10 subgrouping from an isotropic Mardia–Dryden distribution of 150 landmarks in two dimensions (hence our usual count of 300 shape coordinates) based on a template that is a perfect 15 × 10 grid. Each of four different bgPC1–bgPC2 scatters is presented next to the grid for its bgPC1 as graphed by its effect on the template at Procrustes length 1. At upper left is the result of the standard bgPCA dataflow from Section 2: the same fictitious equilateral triangle we have seen many times already. The group separation is perfect, as always, but the corresponding bgPC1 grid is clearly biological nonsense. (Mitteroecker and Bookstein 2011 used this same graphical strategy to mock the analogous vector output from a CVA.) The pair at upper right presents the same bgPCA scatter and bgPC1 grid interpretation after the simulated data set has been deflated by a slope of −0.5, as defined in Bookstein 2015 — this might be the equivalent of mis-analyzing a data set comprised mainly of semilandmarks as if they were landmarks instead, as discussed by Cardini (2019).

The pair of panels at lower left represents the situation after deflation at slope −1.0, the case of self-similarity (the null model in Bookstein 2015). For these self-similar data the bgPCA algorithm continues to pull the group means apart, but the clusters now fail to separate, and after smoothing to this degree of realism the grid seems reminiscent of phylogenetic applications in its offering of diverse potential characters at a diversity of geometric scales. The figure’s final pair of panels, lower right, corresponds to deflation at slope −2, the maximum degree of integration seen in real growth data. Now the basic bgPCA pathology on which this paper centers is almost gone — the degrees of freedom are no longer close enough to spherical for the MPT to have much effect (but you can still see it trying) — and the bgPC1 grid now resembles actual published grids for growth gradients such as were exemplified in Bookstein 2019. In summary, the more integrated the shape variation, the lower the effective number of variables contributing to the MPT formula, the lower the fictitious group separation in bgPCA scatterplots, and the smoother and more realistic the corresponding transformation “factors.”

**Figure 20.**
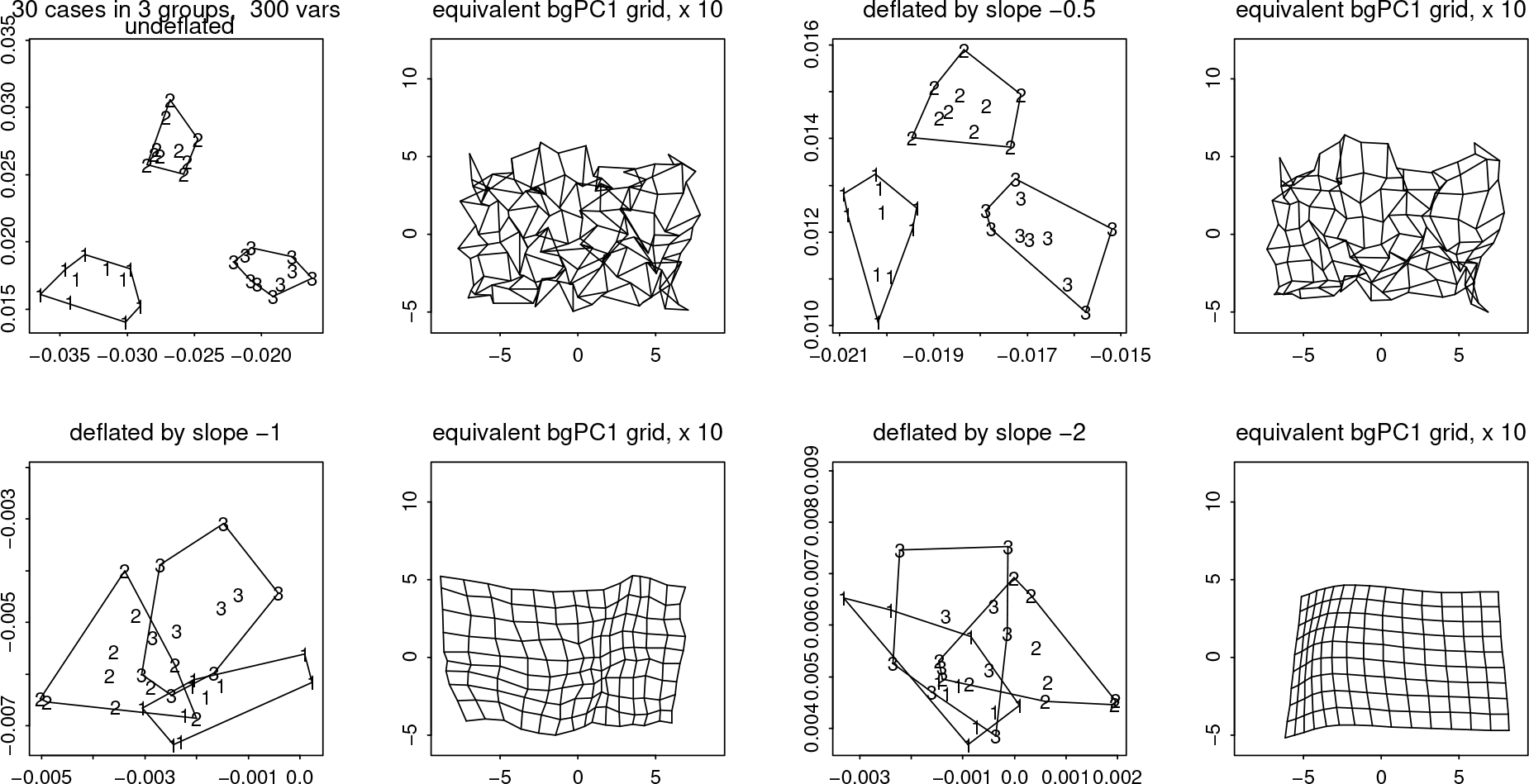
Effect of changing an integration dimension on the fictitious bgPCA plot and the corresponding transformation grid for bgPC1 in a simulated data set of the usual design (30 specimens in three meaningless groups, 300 variables). See text.

In another approach, the low-rank scenario (linear or quadratic terms dominant), the dimensionality of the valid factors present is low enough, and the sum of the variances of the potentially spherical-Gaussian residuals diminutive enough, that the count of true factors contributing to a bgPCA scatterplot is only 1 or 2 regardless of the number of additional dimensions of noise. This variant is illustrated via an evolutionary example (the variation of mammalian skulls) in Figures 7 through 10 of Bookstein (2019).

In terms of the typology of integration laid out in Bookstein (2019), comparisons along a series of deflations with increasingly negative slopes would leave a bgPCA essentially invariant if its effect were at very large scale — a growth-gradient, for example, would be analogous to the single-factor models illustrated in the top row of Figure 19 — but would efface group separations at small scale should they have been fictional after the manner simulated in this isotropic Mardia–Dryden distribution. (Thus the deflation technique suggests itself as an explicit analytic tool for exploring a hypothesis of heterochrony in a high-*p/n* data set.) Deflation greatly attenuates the signal strength of small-scale findings, but so does any other version of PCA; so a search for discriminating “characters” at small scale would not proceed well by any version of today’s GMM, but requires functional or evo-devo arguments instead, followed by the explicit construction of both the loadings and the scores of any hypothesized factors.

The critique of this section can be tersely summarized as follows: **the technique of bgPCA is far from ready for general adoption**, even by disciplinary communities that have been taught tools from the early twentieth century such as principal components analysis that are mathematically welll-characterized already in applications to studies involving far fewer variables than specimens. The bgPCA method has never been subjected to close examination of its remarkably unfortunate tendency to exaggerate or simply invent apparent distinctions under conditions of high variable count, high variation of group size, confounding with within-group factors, and the like. This will be the first in the series of summary mantras that are the gist of the concluding section of this article.

## 5. Concluding observations

Readers should consider Figure 10 very thoughtfully. *That second dimension in the upper panel is not part of the model* — it is wholly an artifact of the bgPCA method. Then the second dimension of the published example below it could itself have been entirely a methodological artifact as well, a software-hallucinated pattern claim unrelated to any evolutionary-biological truth about the genus *Pan*. If a single-factor model on 86 variables for 104 cases can match so closely and saliently a bgPCA of the same structure claiming the existence of two real factors, a bgPCA that I myself co-published as recently as eight years ago, clearly something is drastically wrong with our toolkit for inferences from multivariate analysis of data sets of even higher *p/n* ratio — after all, this was a *p/n* ratio of only 0.83. Likewise the ANOVA model in Figure 12, or its realization as a real object in Figure 13, should be consternating. We should be *very* chary of analyses that, when applied to classically meaningful explanatory structures, give rise to graphics as ludicrous as this match of an interaction-free analysis of variance to an airborne coffee stirrer.

I offer several weapons (or, in what might prove a better metaphor, prophylactics) for our community to wield in defending against this class of problems.

1. **The bgPCA method can no longer be regarded as just another “standard tool” that can be used routinely by nonexperts and justified merely by a rote citation to earlier publications by me or any other tool developer**. There are simply not enough deep theorems about how it operates in today’s typically high-*p/n* settings. Associated with its many pathologies are too many requirements for confirming plots, resampling strategies (including deflation), and appropriate challenges to the inappropriately attractive explanations it routinely proffers without adequate justification. Papers should no longer appear that, like Détroit et al. 2019 or Chen et al. 2019, simply cite Mitteroecker and Bookstein 2011 or some other single earlier article as authority for presentation of one or more standard bgPCA scatterplots without any of the quantitative challenges or alternative interpretations just reviewed in Section 4. No further bgPCA analyses should be published anywhere without being subject to the challenges reviewed in this article, followed by the imprimatur of an appropriate multivariate expert, and the technique should not be used in dissertations or other student work. It is not like the other multivariate techniques that our graduate curricula teach — even most experts do not yet understand what it gets predictably, disastrously wrong.
2. Every published example of a PCA on a data set characterized by more variables than cases must include explicit declarations of the values of *p* and *n* and their ratio. This requirement should be enforced by reviewers and journal editors alike. When a previously published example is cited as an empirically factual supporting argument, the citing sentence should extract the *p/n* ratio from the text being cited and print it alongside the actual citation.
3. Every published example of a bgPCA or the equivalent PLS in a high-*p/n* setting must include challenges to the claimed findings that acknowledge a-priori these MPT-related pathologies. Such challenges must supersede the ordinary casewise crossvalidations or permutations of group assignment, instead extending to the constructions in Sections 4.4 and 4.5.
4. Analyses that find near-equality of the first two or three eigenvalues of a high-*p/n* PCA or bgPCA should be regarded not as evidence for biologically meaningful structure of the corresponding ordinations, but contrariwise as hints of an underlying sphericity (patternlessness) in the data driving the multivariate analysis. This advice is especially important should group centroids (if grouping is the topic) prove to be suspiciously close to equilateral in their ordination space. Conversely, if within-group factors are apparent in the bgPCA output the analysis should be repeated and fictitious sphericity and separations of clusters checked again after these factors have been identified, estimated, and then partialled out of the data.
5. As demonstrated in Figure 17, bgPCA should *never* be used, even by experts, when the range of group sizes is wide or when any of the groups are truly tiny. I would suggest requiring a count of at least 10 for the smallest group before a bgPCA can even be contemplated, let alone published.
6. Generally speaking, ordinations by linear models like PCA, bgPCA, and PLS should be trusted only to the extent that the dimensions they highlight prove to be factors, meaning, biological causes or effects of which the relevant branches of the biosciences were previously aware. Such factors can be expected to persist into the bgPCA plots even though the algorithm was designed to ignore them. Group separations in high*p/n* settings that emerge only after factors emerge, after the manner of Figure 20, should be viewed with particular suspicion. In particular, claims that axes of separation of group averages in GMM analyses are good factors or biological explanations require far stronger, geometrically more specific prior hypotheses than have been exemplified hitherto in our textbooks and our peer-reviewed citation classics, and need to be visualized as thin-plate splines after they have been computed and sample scores scattered.
7. (This comment is specific to the context of GMM.) Any PCA-based technique of mixed landmark/semilandmark analysis needs to be recomputed after restriction to only the Cartesian coordinates perpendicular to the sliding constraints — the coordinates that have not been relaxed by the sliding algorithm. Also, any analysis of an anatomical configuration that combines semilandmarks with a reasonable number of reasonably distributed landmarks needs to be confronted by the parallel analysis that involves only the landmarks, as in Figure 5.73 (page 447) of my 2018 textbook. When the two analyses generate substantially similar ordinations, then the higher the ratio of the larger count of variables to the smaller count, the greater the additional confidence in the inferences that follow from the pair.

I am aware of the consequences that would ensue if this critique of the bgPCA method ends up widely accepted. The trend of increasing evocation of bgPCA in applied papers from whatever disciplinary context would need to be reversed and then drop to nearly zero. A substantial number of peer-reviewed empirical claims that invoke bgPCA as one of their analytic tools, such as two recent well-publicized announcements about human evolution (Détroit et al. 2019 and Chen et al. 2019), may need to be reargued or weakened, and some dissertation chapters abandoned, and also there might need to be a substantial extent of “unlearning” by colleagues who have relied on methods such as permutation testing to assert “statistical significance” for claims of biological significance based in highdimensional pattern analyses like those pilloried here. But such costs must be paid, because our disciplinary community is responsible for having accepted bgPCA into its toolkit of routine methods without a proper vetting.

*We must not continue to be so naïve about the limits of our intuition regarding these high-dimensional “data reduction” techniques*. Let me put this as bluntly as I can: covariance structures on a mere 30 cases in 300 dimensions, or any other high-*p/n* data design, do not resemble scatter ellipses on paper in any manner relevant to empirically valid explanations of organismal form. In these high-dimensional spaces the desideratum of maximizing “explained variance” makes no biological sense, not when we can’t even explain the list of variables with which we’re working, and likewise the notion of valuing more highly the pairs of linear combinations that happen to be uncorrelated; and likewise, too, the strategy of analyzing group means without attending to the structure of multidimensional within-group variability, including the standard error of the mean vector that is the link to the bgPCA arithmetic.

To my knowledge, prior to my textbook of 2018 no vademecum of applied multivariate statistics for biologists, however sophisticated, ever warned its readers about the pitfalls of high-*p/n* PCA as reviewed in Bookstein (2017), of which the bgPCA technique dissected here is a particularly blatant case. It is not unthinkable that PCA itself no longer makes sense in organismal applications now that our data sets can comprise such huge numbers of variables. After all, as Ian Jolliffe (2002:297) wisely noted,

> PCA has very clear objectives, namely finding uncorrelated derived variables that in succession maximize variance. If the [phenomena to be explained] are not expected to maximize variance and/or to be uncorrelated, PCA should not be used to look for them in the first place.

The sciences of organismal form are among those for which neither maximizing variance nor enforcing noncorrelation makes much sense in any megavariate context, whether GMM or another. And Jolliffe’s statement applies as well to bgPCA as to any other application of PCA in the course of analyzing organismal form.^5^

The scenario in Figure 1 is an artifice, but those in Figures 8, 10, 11, and 20 are realistic. In the light of examples like these, why would any applied biometrician of organismal form spend time constructing projections optimizing unrealistically symmetric figures of merit like “explained variance” or “explained covariance,” instead of postulating factor models a-priori, based on established biological theories or replicated experimental findings, and then testing them against data that were newly accumulated for the specific purpose of testing those theories? The symmetries of multivariate statistical method are no match for the subtleties of organismal form as we currently understand its structured variability, its vicissitudes, and its genetic and epigenetic control mechanisms. For a much deeper discussion of these matters, including deconstruction of numerous historical precursors, see my extended commentary in Bookstein (2019). If the examples in Figure 10 or Figure 20 are representative, the only reliable dimensions of a bgPCA analysis are those showing the largest within-group variances, even though that is the information that the bgPCA method was explicitly designed to ignore. Then quite possibly the explicit purpose of bgPCA analysis, the study of group average differences without referring to within-group variability, is self-contradictory in GMM and similar high-*p/n* settings.

## Acknowledgments

Andrea Cardini (University of Modena and University of Western Australia) was the first in our group to notice bgPCA’s clustering pathology, and the idea of a pair of papers scrutinizing applications of bgPCA to high-dimensional GMM data sets, particularly insofar as they ignore the implications of the MPT, was his. Our discussion sharpened over many months of interaction also drawing in Jim Rohlf (Stony Brook University), and Paul O’Higgins (University of York), discussions leading to Cardini et al. (2019) as well as this paper. Norman MacLeod (London and University of Nanjing) guided me through a thoughtful review of the twentieth-century origins of the method criticized here, and Michael Perlman (University of Washington) was a helpful guide into the literature of the MPT at the time all this work began. I am also grateful for the comments of XX anonymous reviewers for this journal. Some aspects of the argument here were acknowledged earlier at the conference “GMAustria19,” Department of Botany, University of Vienna, March 27–28, 2019. As ever I am grateful to Philipp Mitteroecker (University of Vienna), colleague and friend, for arguments and insights both before and after the 2011 joint article that I am here attempting to revise so radically by discrediting the specific technique that was most highly recommended there, and also for his profound critique of an early draft that eventually engendered a major expansion of Section 4.

## Footnotes

1 I am ignoring an intercalated algebraic step of theirs, a standardization of each measured variable by its within-group standard deviation. In GMM applications, standardization is instead by the Procrustes metric that all the shape coordinates share. Incidentally, Yendle and MacFie might also be cited as the first to notice the identity of the PLS and PCA approaches to this computation when, in a closing comment, they noted how their analysis can be implemented using “an algorithm such as NIPALS.” That acronym stands for “nonlinear iterative partial least squares,” the original name put forward by Herman Wold and his son Svante Wold for what was soon to be given a shorter moniker, PLS.

2 I strongly urge all the authors of all 115 of these articles to revisit their multivariate inferences by more thoroughly tested methods capable of checking for the pathologies of the bgPCA approach that their publications originally exploited. The upward trend of these counts makes that advice ever more urgent. Regarding the scientometrics of misinformation in general, see O’Connor and Weatherall, 2018.

3 Current usage, 10^12^, not the former British usage, which meant 10^18^.

4 I am arguing that high-*p/n* divisions of organismal biology like GMM should restrict statistical significance testing to what Paul Meehl (1967) called its “strong version”: tests expected not to contradict but instead to *support* the inference that your null distribution model is at least approximately true. Stated as an aphorism: in the highly multivariate context of good GMM, statistical significance testing should be applied only when the null model is likely to suit your data, exactly or closely enough. Meehl argues that physics exploits the strong version of significance testing, psychology, alas, the weak version only. In its logic of inference, organismal biology ought to resemble physics more than psychology. See in general Bookstein 2014.

5 As Yendle and MacFie (1989) noted in the earliest announcement of the bgPCA method, the factors it produces are uncorrelated in the original data space. Such a claim would seem irrelevant to any actual biological argument. In fairness, it must be noted that the journal in which their article appeared was aimed at chemists, not biologists — possibly chemists *do* prefer their factors to be uncorrelated. It should also be noted that in the first of the two examples of the Yendle–MacFie article, group sizes are identical across the groups, and in their second example the subsample counts vary only between 8 and 12, so the second fatal pathology identified in Section 4.3 does not apply.

